# Evolution of sex-biased gene expression during transitions to separate sexes in the *Silene* genus

**DOI:** 10.1101/2023.10.02.560480

**Authors:** Djivan Prentout, Aline Muyle, Niklaus Zemp, Adil el Filali, Bastien Boussau, Pascal Touzet, Alex Widmer, Jos Käfer, Gabriel A.B. Marais

**Affiliations:** Department of Biological Sciences, Columbia University, New York, USA; Centre d’Ecologie Fonctionnelle et Evolutive (CEFE), University of Montpellier, CNRS, 34 EPHE, IRD, Montpellier, France; Genetic Diversity Centre (GDC), ETH, Zürich, Switzerland; Université de Lyon, Université Lyon 1, CNRS, Laboratoire de Biométrie et Biologie Evolutive UMR 5558, 69622 Villeurbanne, France; CNRS, UMR 8198 – Evo-Eco-Paleo, Univ. Lille, F-59000 Lille, France; Institute of Integrative Biology, ETH Zurich, Zurich, Switzerland; ISEM, Univ Montpellier, CNRS, IRD, Montpellier, France; CIBIO, Centro de Investigação em Biodiversidade e Recursos Genéticos, InBIO, Laboratório Associado, Campus de Vairão, Universidade do Porto, 4485-661 Vairão, Portugal; Departamento de Biologia, Faculdade de Ciências, Universidade do Porto, 4099-002, Porto, Portugal; BIOPOLIS Program in Genomics, Biodiversity and Land Planning, CIBIO, Campus de Vairão, 4485-661 Vairão, Portugal

## Abstract

Sexual dimorphism is widespread among species with separate sexes and its extent is thought to be governed by the differential expression of thousands of genes between males and females (known as Sex-Biased Genes, hereafter SBGs). SBGs have been studied in numerous species, but rarely in a comparative way, which curtails our understanding of their evolution, especially during multiple independent transitions to separate sexes. We sequenced the transcriptomes of nine dioecious species, two gynodioecious species (separate females and hermaphrodites) and two hermaphrodite species from the *Silene* genus. Our dataset provides access to three independent transitions to dioecy (dating from less than 1 Myo to about 11 Myo). We demonstrated that male-biased expression emerges first during a transition to separate sexes, later followed by female-biased genes. Furthermore, we showed that, despite a mixture of selective regimes, positive selection significantly affects the evolution of some SBGs. Overall, this study provides new insights on the causes of SBG evolution during transitions to separate sexes.

**Teaser:** This study describes the evolution of sex-biased gene expression during a transition to separate sexes in plants.

## Introduction

Separate sexes (i.e. gonochorism in animals and dioecy in plants) is the sexual system of 95% of animals and 5% of flowering plants (*1–3*). The differences in the phenotypes of males and females (called sexual dimorphism) can affect the physiology, morphology, and other life history traits (*4–7*). The strength of sexual dimorphism varies widely between species and can be more important than phenotypic differences between individuals of the same sex (*8*). In several species with genetic sex determination, only one or two genes are sufficient to determine the sex of individuals (*9, 10*). The sex determining genes then lead to the activation of a regulatory cascade where both transcription factors and hormones determine the differential expression of up to thousands of genes between males and females. The genes that are differentially expressed between males and females, the so-called Sex-Biased Genes (SBGs), are common in dioecious plants (from 2 to 17% of all expressed genes) and are distributed along the entire genome (*11–14*). Sex-Biased Gene Expression (SBGE) has been extensively described in several animals and to a lesser extent in some plant species (discussed in (*8, 15*)). Previous studies have shown that the proportion of SBGs could vary significantly among tissues and developmental stages (*7, 12, 16*).

Despite numerous analyses of SBGE conducted to date, very few have been done in a comparative way. Therefore, the evolutionary forces at play remain an open question. For example, while a study in birds suggested that sexual selection (approximated by the intensity of sexual dimorphism) had driven the evolution of SBGE (17), converse results were found in cichlid fish (18), in the plant genus Leucadendron (19), and in brown algae (20), where genetic drift was likely to be the strongest evolutionary force driving SBGE. These studies question the common belief that the extent of sexual dimorphism is correlated to the number of SBGs (*8*).

In flowering plants, dioecy has evolved between 871 and 5,000 times independently (*1*), thus providing an exceptional opportunity for comparative analyses. Transitions from hermaphroditism to dioecy are thought to require an intermediate step, often through monoecy (female and male flowers on the same plant) or through gynodioecy (separate female and hermaphrodite individuals) (*2, 10, 17–19*). The latter assumes the invasion of the hermaphrodite population by a male-sterile (female) mutant, leading to gynodioecy (reviewed in (*2*)). Theoretical work suggests that hermaphrodites in gynodioecious populations gain most of their reproductive success through their male function (*17, 20, 21*), the loss of the female function in these individuals can be selected when it increases male fitness, which can lead to the evolution of dioecy. The steps to dioecy through the monoecious pathway have received less attention from modellers so the precise events and the associated selective pressures are less well formalised (*22*). To our knowledge, no comparative study has explored the evolution of SBGE in the monoecy nor the gynodioecy pathway in plants.

The *Silene* genus is a model for studying the evolution of plant sexual systems (*23, 24*). At least three independent transitions to dioecy have been reported in *Silene* (*25–27*). It is likely that these transitions occurred through the gynodioecy pathway, as the genus contains many gynodioecious species. Dioecy evolved ∼11 My ago in the *Melandrium* section, consisting of five dioecious species (*S. latifolia, S. dioica, S. heuffelii, S. marizii and S. diclinis*, (*25, 28–30*)). Dioecy is probably younger in section *Otites* (∼2.3 My; (*27, 31, 32*)) and likely of very recent origin in *S. acaulis* ssp *exscapa* (less than 1 My, (*26*)). XY sex chromosomes share a common origin in the *Melandrium* section (*33*). In the *Otites* section, *S. otites* has ZW sex chromosomes, while *S. pseudotites* and *S. colpophylla* have XY sex chromosomes (*27, 31*). The ZW and XY systems evolved from different autosomes, although the exact evolutionary history (possibly involving introgression or turnover) is not known (*27*). So far, no sex chromosomes have been identified in *S. acaulis*. If a non-recombining region exists in *S. acaulis*, it is likely to be very small and carry only a few genes (*34*).

The repeated independent evolution of separate sexes with different ages of dioecy in *Silene* makes it an interesting model to study SBGE evolution in a comparative framework. So far SBGs have only been studied in *S. latifolia (12*).

In this study, we characterise SBGE in the nine *Silene* dioecious species listed above (from the three independent transitions to dioecy) and two gynodioecious *Silene* species (*S. vulgaris* and *S. nutans*). We use two hermaphrodite outgroup species (*S. paradoxa* and *S. viscosa*) to compare SBGE in dioecious and gynodioecious species to homologous hermaphrodite expression. With this dataset, we aimed to address the following questions: (1) Are there differences in the timing of the evolution of female- and male-biased genes? (2) Are gene expression changes occurring mostly in one sex, as previously suggested in *S. latifolia* (*12, 35*)? (3) Do the same genes repeatedly become sex-biased in the independent transitions towards dioecy? (4) What evolutionary forces shape SBGE evolution, drift or selection?

## Results

### Transcriptome assemblies and mapping results

The *S. nutans* and *S. vulgaris* assemblies were composed of 23,836 genes and 31,526 genes, respectively (see Supplementary Table S1). A BUSCO (version 3.1.0; (*64*)) analysis showed that the *S. vulgaris* transcriptome assembly is more complete than that of *S. nutans* (Supplementary Table S1).

We mapped the RNA-seq data of the 9 dioecious species on both transcriptome assemblies and found that the mapping rate on *S. vulgaris* assembly was more similar among all the species (50%) than the one on *S. nutans* assembly (Supplementary Table S2). Coupled with the BUSCO results, we decided to keep the mapping on *S. vulgaris* for the rest of the analyses.

### Numbers and proportions of SBGs in dioecious and gynodioecious species

We used three different tools to identify sex-biased genes: (1) DESeq2, (2) edgeR, (3) Limma-Voom. These three methods differ slightly in the way they assess sex-biased gene expression due to differences in the family distribution used for read counts. The two firsts rely on a negative binomial distribution and the latter on log-normal distribution. To limit biases, the genes with a log(foldchange)>2 and a *p-*value<10^-4^ in at least 2 of the three methods were considered as sex-biased. The number of sex-biased genes and the number of sex-biased genes identified as autosomal or sex-linked by SEX-DETector are presented in Supplementary Table S3. Figure 1 represents the proportions of sex-biased genes among expressed genes for the eleven species.

**Figure 1:**
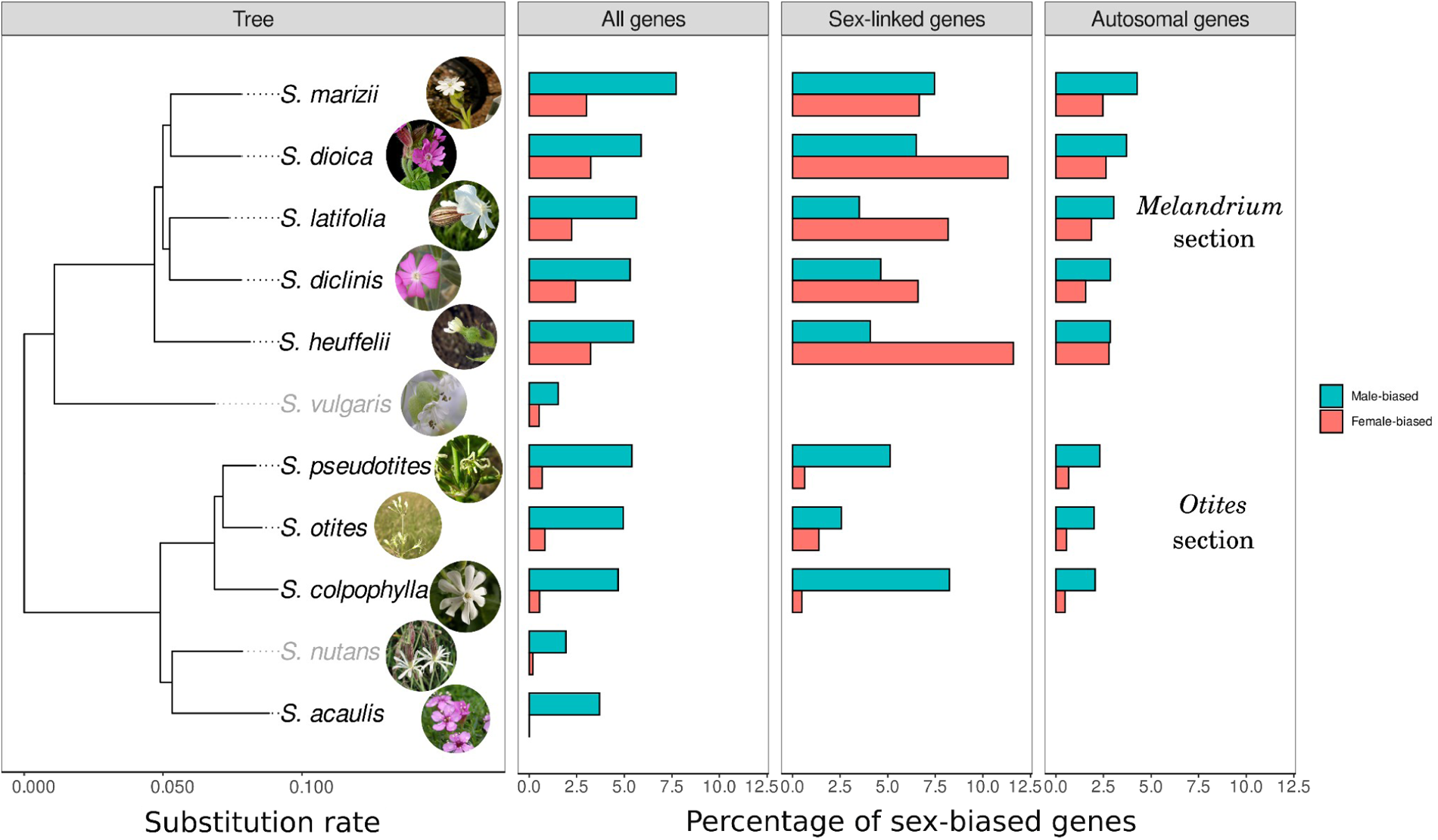
Phylogenetic reconstruction (left panel) of the eleven *Silene* species used in this study and sex-biased gene proportions in each species (the outgroup *Dianthus chinensis* was removed for this figure, see Supplementary Figure S12 for a complete phylogeny). Gynodioecious and dioecious species names are written in grey and black, respectively. The proportion of expressed genes which are female-biased (red) or male-biased (blue) is shown for different gene subsets: all expressed genes (middle left panel), sex-linked genes (middle right panel, only for Otites and Melandrium sections) autosomal genes (right panel, only for Otites and Melandrium sections). See Supplementary Table S3 for detailed gene numbers. (source of the *S. colpophylla* picture: www.earth.com).

For the rest of the analysis, we will report the genes over-expressed in hermaphrodites of the gynodioecious species as male-biased, as hermaphrodites of gynodioecious species reproduce mainly through the male function (*21*). Also, we checked for methodological biases by repeating the analyses with different *p*-values and fold-change thresholds for sex-biased genes inferences, and to test for an effect of sample size we repeated all analyses with 4 males and 4 females only for all species (Supplementary Figure S1 to S5 and Supplementary Tables S4 and S5). We found qualitatively similar results on these controls.

As shown in Figure 1, species in section Melandrium have the highest proportion of female- and male-biased genes. This difference is stronger for female-biased genes, with species in section *Melandrium* having about 4 times more female-biased genes than those in section *Otites*, and ten times more than gynodioecious species. The proportion of male-biased genes is much lower in gynodioecious species than in sections *Melandrium* and *Otites*, with *S. acaulis* being somewhat intermediate. Autosomal and sex-linked genes were identified for the eight dioecious species with a known pair of sex chromosomes. The pattern of SBG proportions is similar between autosomal genes and all genes, as expected as most SBGs are autosomal, with male-biased genes being more numerous than female-biased genes (Figure 1). Strikingly, however, female-biased genes are overrepresented in sex-linked genes compared to male-biased genes in section *Melandrium*, but not in section *Otites*.

### SBGs accumulate over time after the evolution of separate sexes

We used a generalised linear model to investigate the relationship between the number of SBGs and the age of dioecy (using a negative binomial family, and by accounting for the different number of expressed genes among species through adding an offset, see Materials and Methods for details). A significant positive correlation was found between the age of dioecy and the number of female-biased, male-biased and the total number of SBGs (p < 10^-7^, R^2^ > 0.9, Supplementary Table S6, Fig. 2). The number of male-biased genes seemed to reach a plateau as the age of dioecy increased (Figure 2). We therefore used a polynomial regression for the relationship between male-biased genes and the age of dioecy (Materials and Methods equation 1, Supplementary Table S6), which explained more variance compared to a linear model. The same was true for the relationship between the number of total SBGs and the age of dioecy (Materials and Methods equation 1, Figure 2, Supplementary Table S6). On the other hand, the number of female-biased genes significantly increased with the age of dioecy without reaching a plateau (Materials and Methods equation 2, Figure 2, Supplementary Table S6). The positive correlations between the number of SBGs and the age of dioecy remained significant when accounting for species phylogeny using generalised least squares (equation 4, Supplementary Table S6).

**Figure 2:**
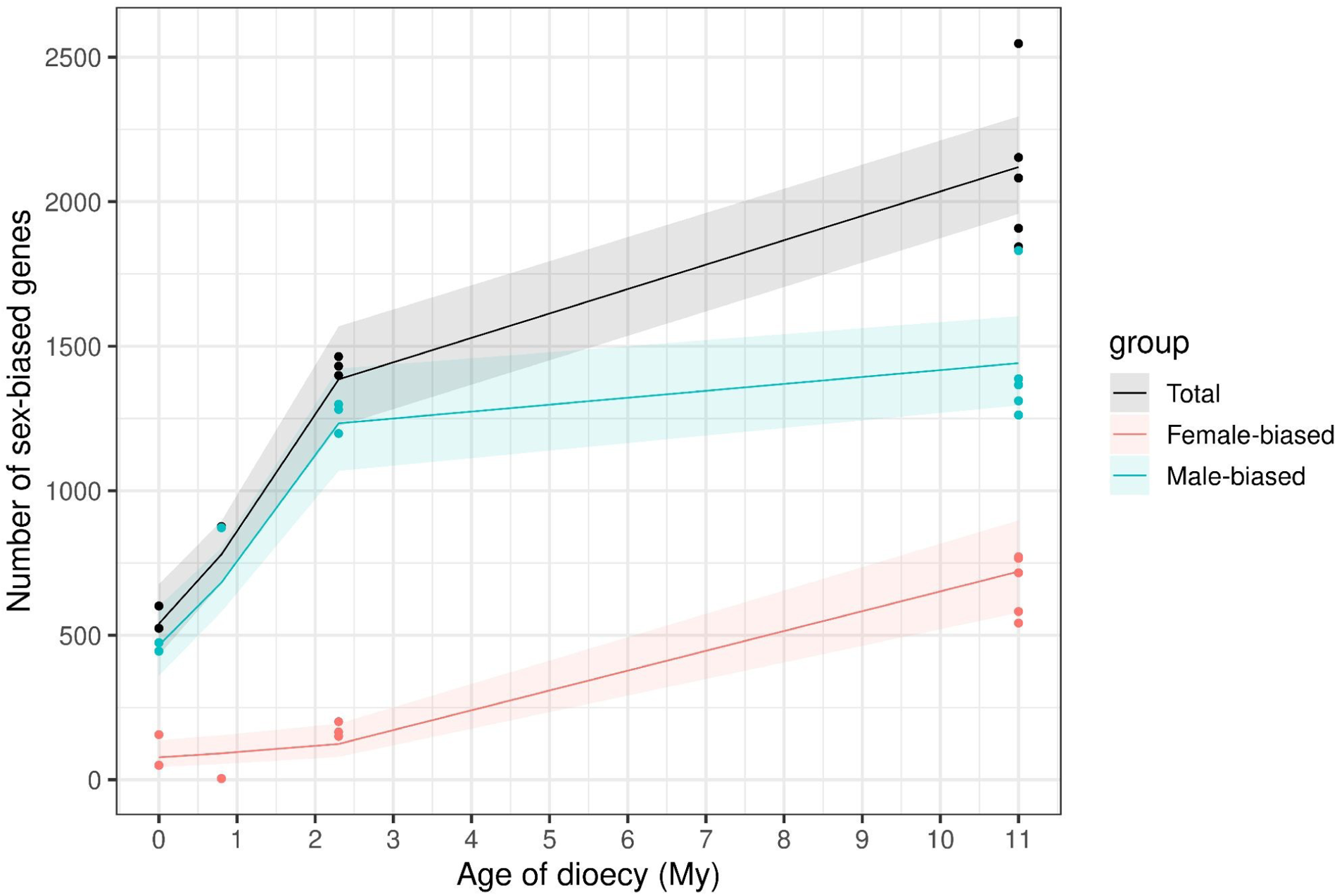
Number of sex biased genes as a function of the age of dioecy (separate sexes) in Million years (My). Gynodioecious species were plotted with age zero of dioecy, *S. acaulis* with age of dioecy 0.8 My, the Otites section 2.3 My and the Melandrium section 11 My. Total sex-biased gene numbers are shown in black, female-biased genes in red and male-biased genes in blue. Dots show the observed data, lines illustrate the predicted values by the generalised linear model detailed in Materials and Methods (equations 1 and 2), ribbons stand for the 95% confidence interval of predicted values. All regressions were significant with p < 1x10^-7^ and R^2^>0.9 (p-values and R^2^ of the models can be found in Supplementary Table S4). For total sex-biased and male-biased genes, the best model included a plateau, specified by a polynomial regression (see Supplementary Table S4 for details). The number of female-biased genes kept increasing with the age of dioecy without reaching a plateau.

The positive correlations observed between the number of SBGs and the age of dioecy suggest it takes time for SBGE to evolve, as species with older dioecy have more SBGE. Male-biased genes evolve early, as they are already present in gynodioecious species, but, after a few My, their numbers reach a plateau in dioecious species. Female-biased genes evolve later, after the transition to dioecy. We did not detect a plateau for female-biased gene numbers over time, but older dioecious species should be studied to address this question.

### High turnover of SBGs in *Silene*

The proportion of species-specific SBGs is larger than that of SBGs shared among several species (Figure 3). Indeed, there is no species for which the number of specific SBGs is smaller than the number of SBGs shared with at least another species. This result shows that there is a high turnover in SBGE evolution in *Silene*. This is consistent with studies in other organisms (*65, 66*).

**Figure 3:**
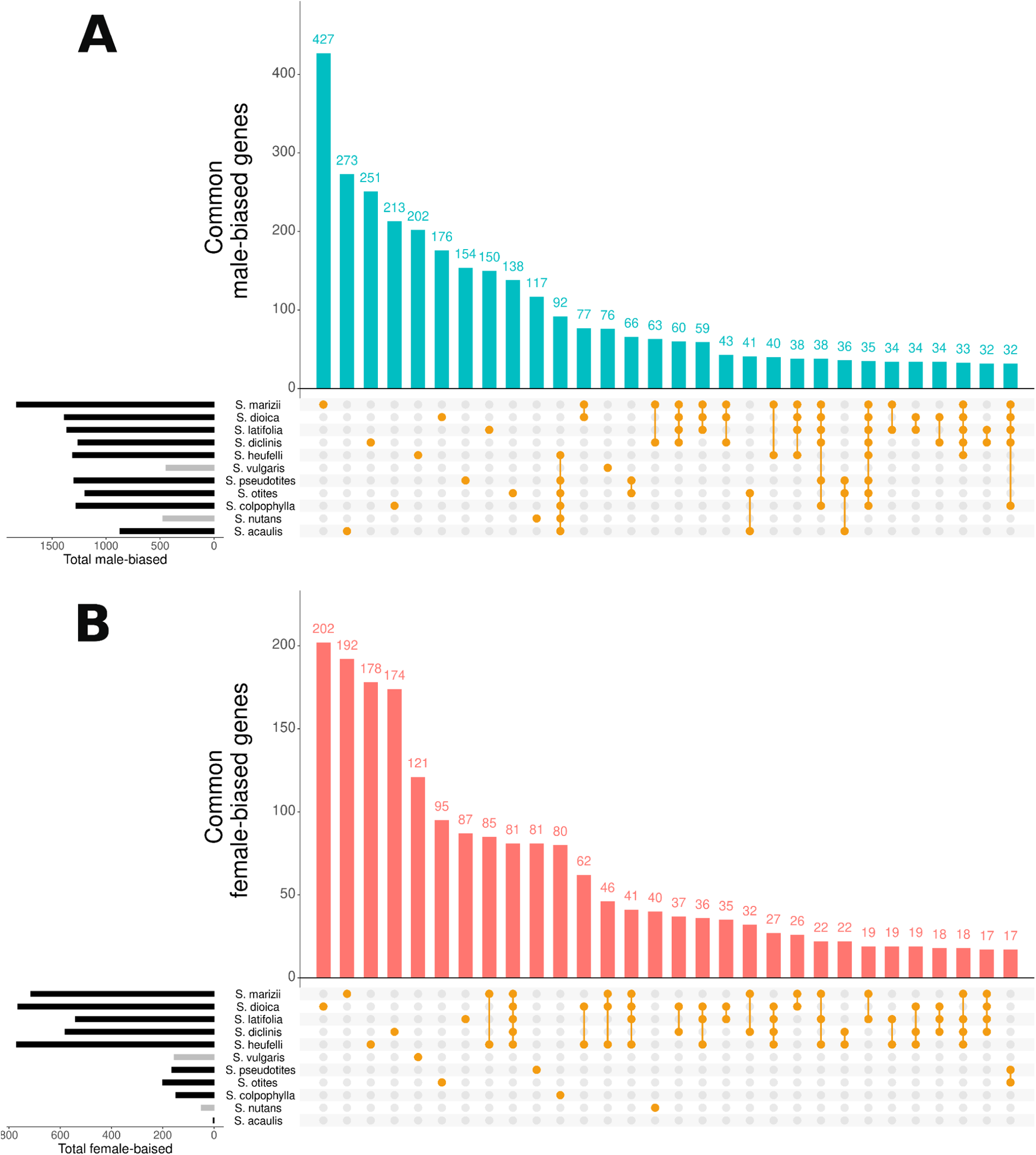
Histogram of male-biased genes (A) and female-biased genes (B) indicating the number of genes that are unique or shared between several species. The yellow dots under each bar indicate the species in which the genes are sex-biased (for example, 192 female-biased genes are unique to *S. marizii* and 85 are shared among *S. marizii* and *S*.*heuffelii*). For both graphs, only the 30 first bars have been represented (see Supplementary Figure S13 & S14 for additional data).

### Selective pressures on SBGE evolution

In order to test whether SBGE evolved under positive selection, we calculated the Δ_X_ statistic for each species and each gene. This statistic summarises the change in expression of a gene in a focal species with respect to an outgroup species, normalised by the standard deviation in the focal species (see Materials and Methods). It is based on the double expectation that positive selection leads to larger than average changes in expression levels between species, as well as more similar expression levels between individuals within a species (*58, 67, 68*). We considered genes with Δ_X_ values higher than the 75 quantile as evolving under positive selection (as in (*12*); see below for a more stringent threshold). In order to test for an enrichment of selection in SBGE, the proportion of SBGs and unbiased genes evolving under selection were compared using a Chi-square test for each species, sex and type of sex-bias (Figure 4, Supplementary Tables S7 and S8).

**Figure 4:**
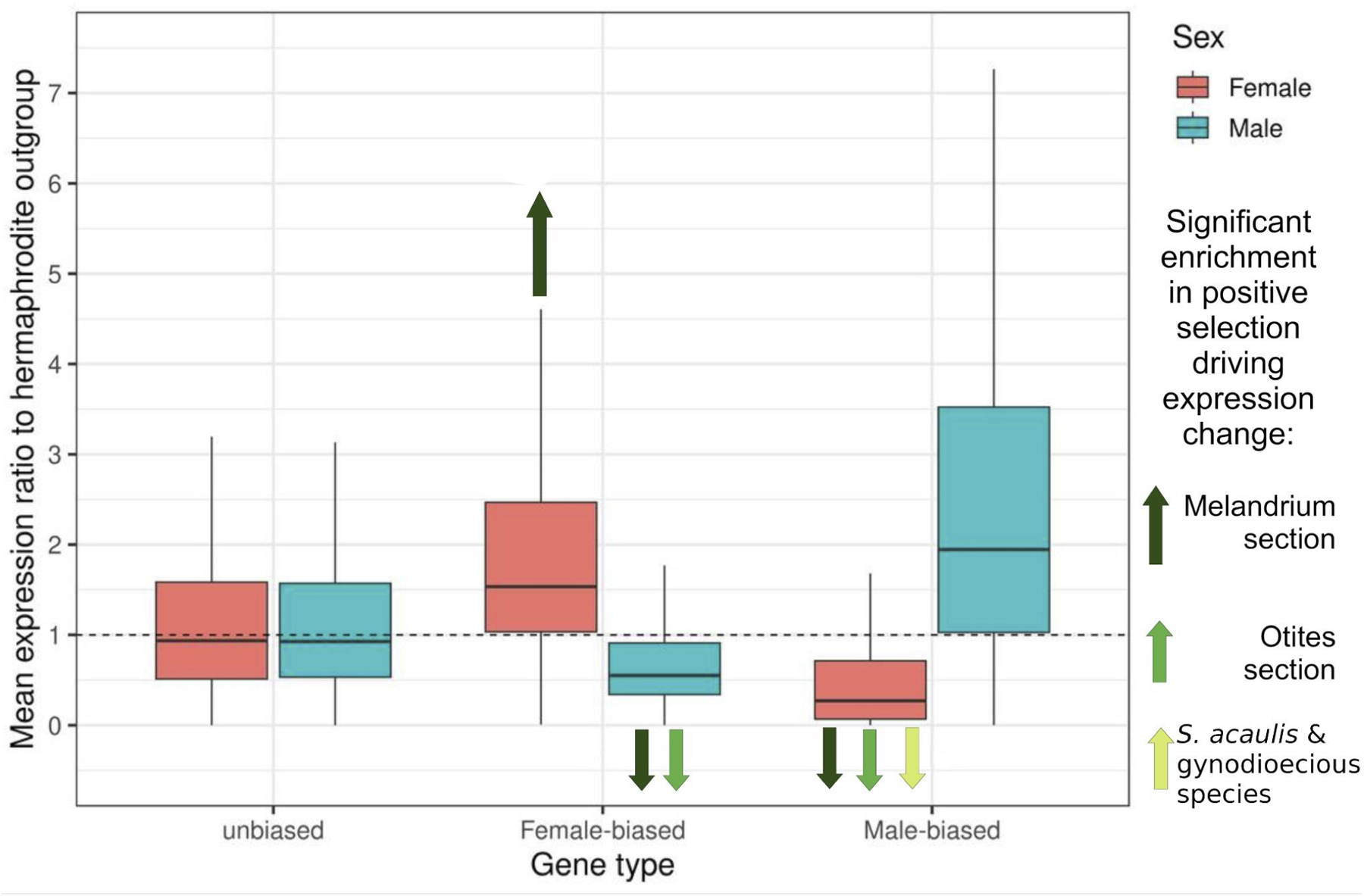
Boxplot of the expression ratio between focal species and their hermaphrodite outgroup. All species were plotted together (see Supplementary Figure S15 for a plot by groups of species). Values higher than one indicate higher expression in the focal dioecious species, and values below one lower expression in the focal species. The arrows summarise the results of the Δ_X_ analysis (Supplementary Tables S7 and S8). Dark-green arrows indicate that sex-biased genes were significantly enriched in selection for increased or decreased expression in four species of the *Melandrium* section. Medium-shaded green arrows indicate that sex-biased genes were significantly enriched in selection for decreased expression in three species of the *Otites* section. The light-green arrow indicates that male-biased genes were significantly enriched in selection for decreased female expression in *S. acaulis* and in gynodioecious species.

As shown in Figure 4, female-biased expression is due both to increased female expression and decreased male expression compared to the hermaphrodite outgroup. Similarly, male-biased gene expression is due both to decreased expression in females and increased expression in males compared to the hermaphrodite outgroup. In order to assess whether these expression changes in SBGs were driven by selection, we tested for an enrichment in selection in SBGs compared to unbiased genes (Supplementary Tables S7 and S8). First, in both sections *Melandrium* and *Otites*, female-biased gene expression that evolved through decreased expression in males compared to the hermaphrodite outgroup was enriched in positive selection compared to unbiased genes. Second, for the *Melandrium* section only, increased expression in females for the female-biased genes is also enriched in positive selection compared to unbiased genes. For male-biased gene expression, significant enrichment in positive selection was also found among genes for which expression decreased in females compared to the hermaphrodite outgroup. However, increased male expression compared to hermaphrodite outgroups was significantly depleted in positive selection for male-biased genes in eight out of eleven species, suggesting that male expression evolves mostly under drift or purifying selection in male-biased genes. To differentiate between drift and purifying selection, we considered expression variation within species (standard deviation). Male expression was the most variable in male-biased genes, especially when male expression increased compared to the outgroup (Supplementary Figure S6), suggesting drift was the driving force for the evolution of expression in this category.

Results were qualitatively similar when considering contigs with Δ_X_ > 10 as under positive selection (as done in Scharmann *et al*. 2021 (*69*)), instead of using the quantile 75 (Supplementary Table S9). Using a threshold of Δ_X_ > 10 is more stringent to infer positive selection because the species quantile 75 of the Δ_X_ ranged from 2.5 to 4.6. Results were also unaffected when we repeated the analyses on autosomal genes only (for species without sex chromosomes all genes were kept, for species with sex chromosomes, only genes inferred as autosomal by SEX-DETector were kept, Supplementary Table S10). As another control, we ran the same analysis on leaf tissues (instead of flower buds) in *S. latifolia* only, using *S. viscosa* leaf data as a hermaphrodite outgroup to compute the Δ_X_. We detected a significant enrichment in selection for leaf male-biased genes in *S. latifolia* when male expression increased compared to the outgroup, as well as when female expression decreased compared to the outgroup (Supplementary Table S11).

### Selected feminization of the X chromosome

We investigated whether the increase in the proportion of female-biased genes among sex-linked genes in the *Melandrium* section can be attributed to the feminization of the X chromosome. Feminization of the X is an enrichment of female-beneficial genes on the X, expected to occur because the X chromosome spends two thirds of its time in females and only one third of its time in males (*70, 71*). To test for X feminization, we compared the proportion of female-biased genes on the X that have an expression evolving under positive selection (1) to the proportion of male-biased genes on the X with an expression evolving under positive selection, and (2) to the proportion of female-biased genes on the autosomes with an expression evolving under positive selection. We found that the proportion of female-biased genes evolving under positive selection was significantly higher on the X compared to the other two categories in the *Melandrium* section (Chi-square test; see Supplementary Tables S12 and S13). This shows that the X is significantly enriched in female-biased genes under positive selection in the Melandrium section, suggesting active X feminization. Interestingly, we observed more positive selection signatures for a reduction of expression in males than for an increase of expression in females. These results tend to confirm a feminization of the X chromosomes in the *Melandrium* section. The numbers of female-biased genes in the *Otites* section were too small to conduct such an analysis.

### Evolution of SBGs under drift

We hypothesised that if drift is a strong driver of sex-biased gene expression, species with lower effective population size may have more SBG. On the contrary, if selection is mostly driving the evolution of SBGs, species with higher effective population size (*N*_*e*_) should have more SBGs. In order to test for an effect of the intensity of selection on the number of sex-biased genes, we ran a generalised linear model between the number of sex-biased genes and the synonymous nucleotide diversity *π*_*S*_ (a proxy for *N*_*e*_, equations 3 and 5). No significant correlation was found in either direction (Supplementary Figure S7, Supplementary Table S14), suggesting that drift and selection do not explain differences in SBG numbers among species, or cancel each other.

As done by Scharmann *et al*. (2021), we tested whether sex-biased genes had an increased expression evolutionary rate compared to unbiased genes. To avoid rates being affected by SBGE evolution, we computed rates of SBGE evolution after removing the species for which the gene is sex-biased, so that the rates correspond to expression changes before the gene became sex-biased. We therefore compared expression evolution rates between genes that are always unbiased in all species and genes that are sex-biased in at least one species, after removing species with sex-biased expression from the analysis. We found, as Scharmann *et al*. (2021), that genes that are sex-biased in at least one species have a higher rate of expression change, measured as the mean of absolute PICs (91.91 on average) compared to genes that are always unbiased (51.93 on average, one-sided permutation test p-value < 2.2x10^-16^, Supplementary Figure S8). Scharmann *et al*. (2021) interpreted this result as an indication that genes that become sex-biased have faster-evolving expression levels, even before becoming sex-biased (*69*). To further test this interpretation, we splitted the analysis of PICs for genes that were detected as evolving under positive selection in at least one species and genes that never evolve under positive selection (using the previous Δ_X_ analysis). We found that SBGs that evolve under positive selection in at least one species have the highest rate of expression change, even before becoming sex-biased (94.96 on average, permutation tests p-values < 10^-3^, Supplementary Figure S8). The rate of expression change was lowest for unbiased genes that never evolve under positive selection (mean 10.94, permutation tests p-values < 10^-13^, Supplementary Figure S9). Therefore both positive selection and drift seem to accelerate expression evolutionary rates.

### Functional analysis of SBGs

We explored if the SBGs are enriched for functions or pathways linked to sexual reproduction. We tested the enrichment for four sets of genes (1) the whole set of male- or female-biased genes (2) the male- or female-biased genes in gynodioecious species (3) the male- or female-biased genes in dioecious species (4) the male- or female-biased genes under positive selection (using the previous Δ_X_ analysis, see Supplementary Table S15 for more information). We then looked for enrichment in functions explicitly linked to sexual selection in each of these sets of genes. We also produced wordcloud figures to highlight the main functions or pathways (Supplementary Figure S10 & S11). Only two GO terms are explicitly linked to a reproductive function (*GO:0090567: reproductive shoot system development* and *GO:0048437: floral organ development*). Those two GO terms are found while testing the whole set of female-biased genes against the whole annotation.

## Discussion

We studied Sex-Biased Gene Expression (SBGE) in the flower buds of eleven sexually dimorphic *Silene* species, including nine dioecious species and two gynodioecious species. The nine dioecious species originated from three independent transitions to dioecy. The youngest transition occurred less than one million years ago and the oldest approximately eleven million years ago (*26, 28, 29*).

Overall, the 11 species displayed more male-biased genes than female-biased genes (Figure 1), which has already been observed in various plant species before (*15*). Two possible explanations for this observation are (i) that drift is stronger in males due to a more variable reproductive success and a smaller effective population size, or (ii) stronger sexual selection in males due to strong competition among males. The age of the dioecy is positively correlated with both the number of male-biased genes and the number of female-biased genes. The dynamics, however, seem to differ between sexes: male-biased gene numbers increase early and seem to reach a plateau, whereas female-biased genes evolve more gradually and continuously. A possible explanation for the plateau reached by the number of male-biased genes is that an equilibrium is attained among mutations, selection and drift. Mutations create new SBGs in dioecious species and selection and drift filter them through time.

In gynodioecious species, we consider genes with higher expression in the hermaphrodite individuals as male-biased, since the hermaphrodites’ main reproductive output is through the male function (*20*). A possible explanation as to why male-biased genes are already present in gynodioecious species but female-biased genes are nearly absent is that a gene could become male-biased simply through the reduction of expression in female flowers, as a result of the loss of the male function. Therefore, although female flowers evolve first in the gynodioecy pathway, female-biased genes mainly evolve later, when female functions are suppressed in male flowers (i.e., when full dioecy evolves).

The patterns of male- and female-biased genes enrichment are reversed in the non-recombining region of the sex chromosomes in the Melandrium section (species from the oldest transition to dioecy, Figure 1). We tested whether the female-biased genes on the X chromosome bear footprints of positive selection, using the Δ_X_ statistic (see (*58*)). In brief, if selection drove the evolution of a SBG, we should observe a strong change in the expression level and a small variance between individuals of the same sex in a species. The Δ_X_ analysis showed that female-biased genes in the non-recombining region of the X chromosome are significantly enriched in positive selection compared to male-biased genes on the non-recombining region of the X chromosome or female-biased genes on autosomes. This result supports an active feminization of the X chromosome in the Melandrium section.

We used the Δ_X_ statistic to study the selective regime of all the SGBs in the different species. With the exception of female-biased genes in *S. acaulis* (the species from the most recent transition to dioecy and very few female-biased genes), the reduction of expression in males for female-biased genes, or reduction of expression in females for male-biased genes is enriched in genes with an expression evolving under positive selection for all the dioecious species. In contrast, the increase of expression is enriched in genes evolving under positive selection only in females of the Melandrium section for the female-biased genes. This suggests that, even if SBGs result from changes in both sexes, increases in gene expression do generally not occur through positive selection, while the reduction in expression generally does. For gynodioecious species and *S. acaulis* (which evolved dioecy recently), the decrease in female expression of male-biased genes was significantly enriched in selection (Figure 4). These footprints of positive selection differ from a previous analysis in the *Leucadendron* genus (*69*), where sex-biased genes were not enriched in positive selection on expression levels compared to unbiased genes, suggesting that they mostly evolved under relaxed selection. In order to test whether this difference between the two studies was due to the sampled tissue (flower buds in *Silene* versus leaves in *Leucadendron*), we used leaf tissue available in *S. latifolia* and *S. viscosa* to compute the Δ_X_ in *S. latifolia* leaves (Supplementary Table S11). Leaf male-biased genes are significantly enriched in adaptive evolution compared to unbiased genes in *S. latifolia*. Therefore, the differences between *Silene* and *Leucadendron* are not attributable to the sampled tissue.

Some SBG categories are significantly enriched in positive selection (Figure 4), but other categories seem mostly driven by drift as they exhibit a high within-species variation in expression levels (Supplementary Figure S6). We therefore clearly observe the effects of both selection and drift on SBGE evolution in *Silene*. This is also visible when studying expression evolutionary rates (Supplementary Figure S9). In our *Silene* dataset, sex-biased genes that are never under selection in any species (according to the Δ_X_ analysis) have faster expression evolutionary rates than unbiased genes that are never under selection, suggesting that SBGE leads to faster expression evolutionary rates because of drift. Expression evolutionary rates are even further accelerated by selection, because SBGs that evolve under selection (according to the Δ_X_ analysis) in at least one species have higher expression evolutionary rates than SBGs that never evolve under selection (Supplementary Figure S9). We therefore confirm the theoretical prediction by Dapper and Wade (*72*) that SBGs mainly evolve under relaxed selection, with some exceptions. For some SBGs, positive selection is especially strong and contributes to accelerated rates of expression evolution in *Silene*.

To our knowledge, this analysis is the first comparative analysis of SBGE between gynodioecious species and dioecious species from independent transitions to dioecy. Indeed, despite numerous analyses of SBGE conducted to date, very few have been done in a comparative way (but see (*65, 66, 69, 73*)). This limits our understanding of its evolution and the evolutionary forces that shape it, especially in a transition from hermaphroditism to dioecy or gonochorism. Here, we show that the proportion of sex-biased genes correlates with the age of dioecy and that, through the gynodioecious pathway, male-biased genes emerge first. Also, our results support a combined action of positive selection and genetic drift in the evolution of SBGE. This nuances the theoretical prediction that most sex-biased genes should evolve under relaxed selection, simply because sex-specific expression reduces selection coefficient of a gene (*72*). Our study therefore calls for more investigation in other groups to enlighten why in some species SBGE seems to evolve mostly under drift (like *Leucadendron*), while other species show positive selection driving SBGE (such as *Silene*).

## Materials and Methods

### Data

#### Crossing and sequencing for the *Melandrium* section

For *S. latifolia* and *S. vulgaris*, data have been reused from Zemp *et al*, (2016) and Zemp *et al*. (2018)(*12*),37)). For *S. dioica, S. heuffelii, S. marizii and S. diclinis*, we have crossed a female and a male from two different populations. Seeds from the crosses were sown to produce F1 individuals. Three flower buds (∼2-3 mm) without the calyx were sampled for multiple females and males and their parents (see Supplementary Table S16 for details on sample sizes). High quality RNA (RIN > 9) has been extracted using the Plant Total RNA Mini Kit from Geneaid. RNAseq libraries were prepared using Ultra II RNA Library Prep Kit for Illumina from NEBNext. The libraries were then sequenced on a Hiseq4000 instrument using the paired end 150bp protocol at the Functional Genomic Center, Zurich.

#### Sampling and sequencing of *S. acaulis*

Flower buds from males and females of *Silene acaulis* have been sampled from a natural population, in 2018, at “Rocher Blanc” (“massif d’Allevard”, France, between 2800m and 2900m above sea level).

The RNA preparation was done by the AGAP institute (Montpellier, France). For all individuals, fresh tissues were sent, then flash-frozen (Freshfreeze method). RNA was extracted following the SIGMA protocol and sequenced using the HiSeq3000-HWI-J00115 technology, producing 150bp paired reads.

#### Published dataset for Otites section

For *S. otites* and *S. pseudotites*, we used publicly available data from Martin *et al*. (2019) (*31*). For *S. colpophylla*, we used data from Balounova *et al*. (2019)(*27*).

#### Published dataset for hermaphrodite species, gynodioecious species, and *Dianthus chinensis*

For *S. nutans, S. paradoxa* and *S. viscosa* (the two hermaphrodite outgroups), as well as *Dianthus chinensis* we used publicly available RNAseq data from Muyle *et al*. (2021)(*36*) .

### Transcriptome assemblies

We assembled a *de novo* transcriptome for both gynodioecious species: *S. vulgaris* and *S. nutans*. To this end, we used the tool DRAP (version 1.92,(*37*)), which allows merging and compacting several transcriptome assemblies into a single “meta-assembly”.

For *S. vulgaris*, we first independently assembled two transcriptomes from two hermaphrodite individuals and one transcriptome from a female (tool RunDrap from Drap – default parameters). Then, we merged these three transcriptomes into a single one with RunMeta (from DRAP, default parameters).

For *S. nutans*, we independently assembled two hermaphrodite transcriptomes and two female transcriptomes that we merged in a single one (using the default parameters for RunDrap and RunMeta, as we did for *S. vulgaris*).

DRAP provides several assemblies. We used the one filtered with TransDecoder (https://github.com/TransDecoder/TransDecoder) in order to obtain the Open Reading Frame (ORF) of every gene.

### Trimming, filtering and mapping of reads

We filtered PCR duplicates with Condetri filterPCRdupl(*38*). The reads with a Phred quality score lower than 64 were filtered with trimmomatic (option PE -phred64)(version 0.39; Bolger, Lohse, and Usadel 2014) and the adapters were removed using prinseq-lite (options -trim_tail_left 5 -trim_tail_left 5)(version 0.20.4; (*39*)). Once filtered, the reads were mapped on both transcriptome assemblies using GSNAP (option m 0.1)(version 2019-09-12 ;(*40*)). In order to increase the amount of reads mapped for distant species, we re-ran GSNAP with the SNP-tolerant option and SNPs identified in the first mapping (method previously employed in Prentout *et al*. (2020)(*14*)). This iteration has been done once for each species, except when mapping *S. vulgaris* and *S. nutans* on their respective transcriptome. Samtools version 1.9 (*41*) was used to remove unmapped reads and sort the mapping outputs to bam files (view -F 4 | sort).

### Transcriptome annotation

We annotated the transcriptome of *S. vulgaris* as it is the one that we used for the rest of the analysis (see results section). We used diamond to blast the nucleotide CDS, provided by DRAP, against the nr database (nr version of April 2022)(diamond v2.0.4.142, blastx -p 4 --max-hsps 3 -e 1e-5 -f 5)(*42*). The diamond output (xml format) was provided to Blast2GO for the gene ontology mapping (Blast2GO Basic v6.0)(*43, 44*).

To statistically test if a set of genes is enriched for specific functional pathways, we used two tools: GOstats (version 2.62.0, (*45*)) and clusterprofiler (version 3.15.0, (*46*)), and kept the GO term with an adjusted *p*-value < 0.05.

### Sex-linked genes identification

Sex-linked genes are genes located in the non-recombining region of the Y (or W) chromosome or the homologous region on the X (or the Z) chromosome. In order to identify sex-linked genes in dioecious species, we first inferred the genotype of each individual with Reads2snp version 2.0 (*47*). We accounted for allelic expression bias, without filtering for paralogous SNPs and retained positions with a minimum coverage of 3 (-aeb -par 0 -min 3).

SEX-DETector (*48*), a tool based on the analyses of allele segregation within a cross, was used to identify sex-linked transcripts in every dioecious species of the Otites and Melandrium sections. This tool is based on an SEM algorithm and computes, for each gene, a posterior probability of being autosomal (P_A_), XY (P_XY_) or X-hemizygous (P_X-hemi_). We considered a gene as sex-linked (or autosomal) if P_XY_ + P_X-hemi_ > 0.8 (respectively P_A_ > 0.8), and if at least one SNP was classified as sex-linked (or autosomal) without genotyping error.

Three models have been implemented in SEX-DETector: (1) XY sex chromosome system, (2) ZW sex chromosome system, (3) a system without sex chromosomes.

In all species, the type of sex chromosome has already been identify, so we run SEX-DETector with the corresponding model (i.e. ZW for *S. otites* and XY for the seven other species; (*27, 28, 31*)).

As mentioned in the introduction, no sex chromosomes have been identified in *S. acaulis* so far (*34*). Because their detection is beyond the scope of this analysis, we didn’t detect sex-linked genes in this species.

### Gene expression level and sex-biased gene identification

The expression level of each gene was computed with samtools idxstats (Version 1.9; (*41*)). Then, three tools were used to classify a gene as sex-biased: EdgeR (Version 3.36.0; (*49*)), DESeq2 (Version 1.34.0; (*50*)) and Limma-Voom (Version 3.50.3; (*51*)). We classified a gene as sex-biased if at least two of the three methods classified it as sex-biased with *p-*value lower than 10^-4^ and greater than 2 (log_2_FC > 1).

To infer in which sex the expression changed compared to the ancestral expression level before separate sexes evolution, we computed the mean fragment per kilobase of transcript per million mapped reads (FPKM) for each gene in every individual, and then, the mean FPKM for each gene in each sex (males, females or hermaphrodites). The difference in mean expression between the focal dioecious (or gynodioecious) species and the closest fully hermaphrodite species (either *S. paradoxa* or *S. viscosa*) was used to infer an increase or a decrease in gene expression compared to ancestral expression before separate sexes evolution. For *S. diclinis, S. dioica, S. heuffelii, S. latifolia, S. marizii* and *S. vulgaris*, we used the hermaphrodite species *S. viscosa* as an outgroup. For *S. acaulis, S. colpophylla, S. nutans, S. otites* and *S. pseudotites*, we used the hermaphrodite species *S. paradoxa* as an outgroup.

### Phylogenetic reconstruction

*Dianthus chinensis* was used as the outgroup species for the phylogenetic reconstruction (we used 2 hermaphrodite individuals). In order to build the species tree, we kept transcripts of autosomal genes for which the sequence was available in all individuals of all species (N_individuals_ = 143) with a maximum of 30% of Ns in the sequence (740 genes in total). In each individual, only one of the two alleles of each gene was kept for phylogenetic reconstruction. We concatenated the sequences of the 740 genes in each of the 143 individuals, which led to a sequence of 873,747 bp per individual. The phylogeny was inferred with IQ-TREE2 (version 2.1.3; (*52*)) using the GTR+G4 model and 100 replicates for the bootstraps.

### Linear regressions between the number of sex-biased genes and the age of dioecy

The number of sex biased genes was correlated to the age of dioecy using a generalised linear model with negative binomial 1 family (appropriate for large count data) implemented in the glmmTMB R package (*53*). In order to account for the different number of sampled genes among species, the total number of expressed genes was log transformed and used as an offset. For the total number of sex-biased genes and the number of male-biased genes, the model that explained the most variance was a raw polynomial of degree 2 (equation 1). For female-biased genes, a simpler model without polynomial effects was used (equation 2). Fit of the model to the data was checked using the DHARMa R package by looking at residual plots (*54*).

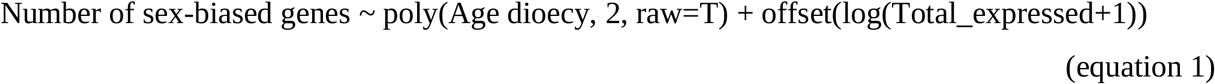

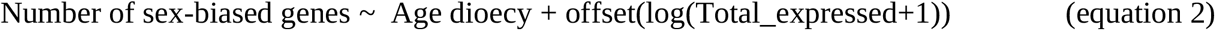

The regression output was plotted using the R package ggeffects to compute predicted values of the model with function ggpredict (*55*). The R^2^ was computed using the function r.squaredGLMM of the R package MuMIn (*56*).

A similar model was run to test for an effect of the effective population size *N*_*e*_ (the synonymous nucleotide diversity *π*_*S*_ was used as a proxy for *N*_*e*_). The negative binomial distribution was used and an offset was included:

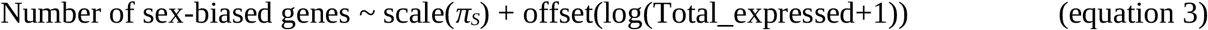

We also ran models that corrected for phylogenetic relationships among species using generalized least squares implemented in the nlme R package (*57*), using function gls and Martin’s correlation structure. Since an offset cannot be implemented in such models, we did not include it. As the negative binomial family is not available is gls, we log transformed the number of sex-biased genes:

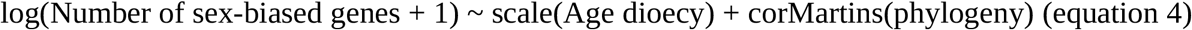

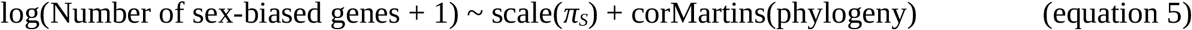

### Δ_X_ analysis

The Δ_X_ has become a widely used statistic in transcriptomics to evaluate selection pressures on gene expression (*58*). In order to compute the Δ_X_ for each sex, gene and species, the expression level of each gene and individual was first determined. The number of mapped reads was computed for each gene and individual using samtools version 1.10 idxstats and was normalised to reads per kilobase per million mapped reads (RPKM) for each contig and individual separately as follows:

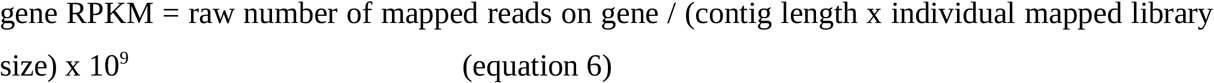

Then, the Δ_X_ was computed for each species and sex separately (i.e. for males and females separately) as follows:

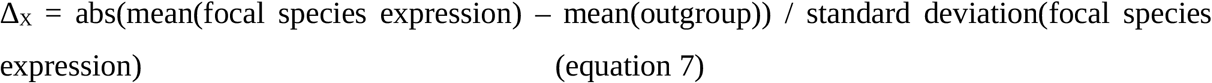

Where abs stands for absolute value. The hermaphrodite species used as the outgroup are the same as in the “Gene expression level and sex-biased gene identification” section.

For each species, contigs with Δ_X_ values higher than the 75 quantile of Δ_X_ values were considered as under positive selection for expression evolution (as in (*12*)). We compared the proportion of SBGs and unbiased genes under selection using a Chi-square test and corrected for multiple tests using Benjamini & Hochberg correction (*59*).

Genes with sex-specific expression were excluded from Δ_X_ analyses, to avoid the confounding effect of including genes that are encoding tissues or functions sex-specific (*i*.*e*. androecium, gynoecium, etc), without the need to involve selection.

We also run this analysis on leaf tissues from *S. latifolia* and *S. viscosa*, to test if different tissues show different results.

### Phylogenetically independent contrasts (PICs)

Phylogenetic independent contrasts (PICs) correspond to the amount of change (here change in expression level) between two taxa divided by the branch length separating them. The mean of the absolute standardised PICs (*60*) per gene was employed as a measure of the rate of expression evolution (*61*). For each gene, PICs were calculated based on the species tree and the mean RPKM expression values per species, using the pic function in APE (*62*). Species exhibiting sex-biased expressions were excluded when calculating PICs, so that the PICs only measure gene expression variation without sex bias (i.e. before the gene became sex-biased). Genes that are sex-biased in at least one species and genes that are always unbiased were then compared for their mean absolute PICs on the basis of 10,000 permutations, using the function permTS in the R package perm v1.0 (*63*).

## Acknowledgments

We would like to thank Theo Tricou for his advice with the phylogenetic reconstruction. This work was performed using the computing facilities of the CC LBBE/PRABI and of the Institut Français de Bioinformatique.

## Author contributions

Conceptualization: GABM, AW

Funding acquisition: GABM, AW

Project administration: GABM, AW

Software: DP, AM, BB

Formal analysis: DP, AM, AeF, BB

Investigation: DP, AM, NZ, BB, PT, AW, JK, GABM

Resources: NZ, AW, PT

Visualisation: DP, AM

Supervision: GABM, JK, AW

Writing—original draft: DP, AM, JK, GABM

Writing—review & editing: All authors

## Data and materials availability

Raw reads are available in the NCBI under the project accession number SUB13729399.

