## Supplementary Material for "Evolution of sex-biased gene expression during transitions to separate sexes in the *Silene* genus"

Djivan Prentout *et al.*

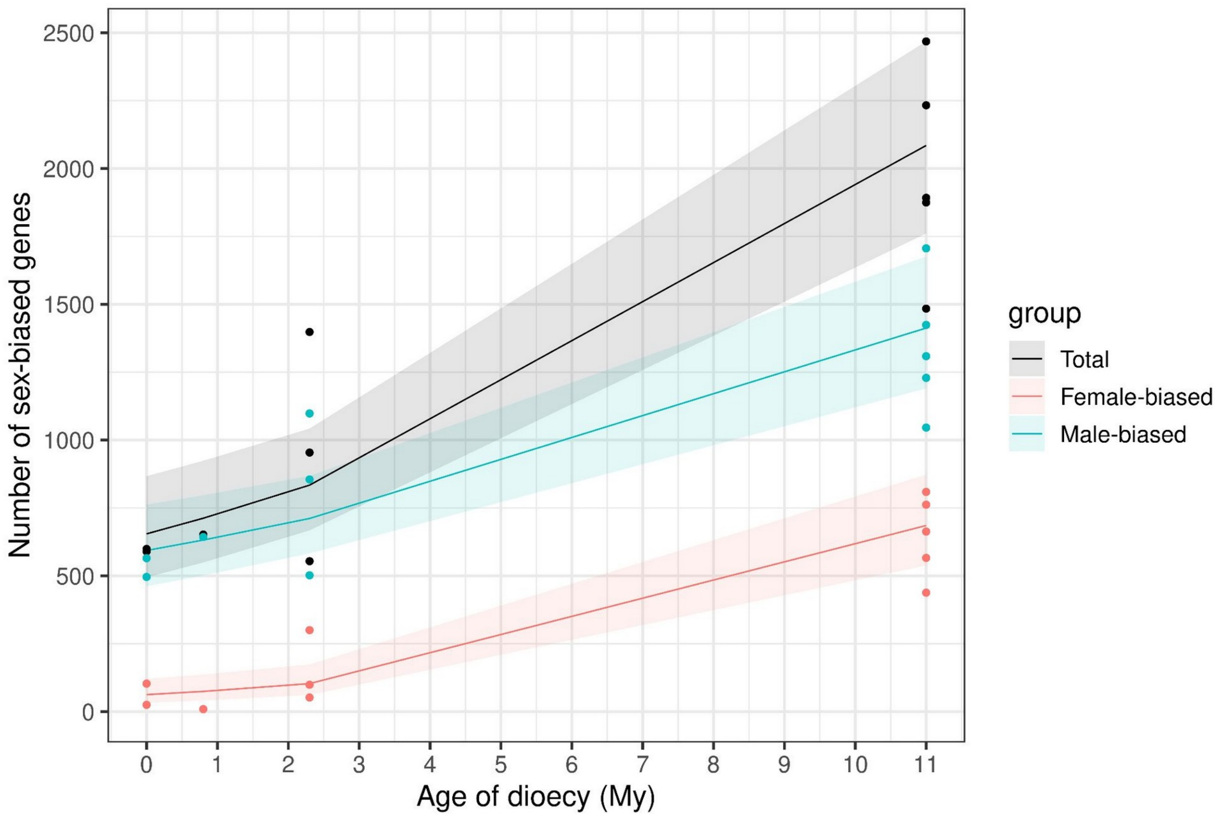

**<u>Supplementary Figure S1:</u>** <u>Same as Figure 2 but with sex-biased genes inferred on a subset of the data with four males and four females for each species.</u> Number of sex biased genes as a function of the age of dioecy (separate sexes) in Million years (My). Gynodioecious species were plotted with age zero of dioecy, *S. acaulis* with age of dioecy 0.8 My, the Otites section 2.3 My and the Melandrium section 11 My. Total sex-biased genes numbers are shown in black, female-biased genes in red and male-biased genes in blue. Dots show the observed data, lines illustrate the predicted values by the generalized linear model detailed in Materials and Methods (equation 2), ribbons stand for the 95% confidence interval of predicted values. All regressions were significant with $p < 1x10^{-7}$ and $R^2>0.75$ (p-values and $R^2$ of the models can be found in Supplementary Table S14).

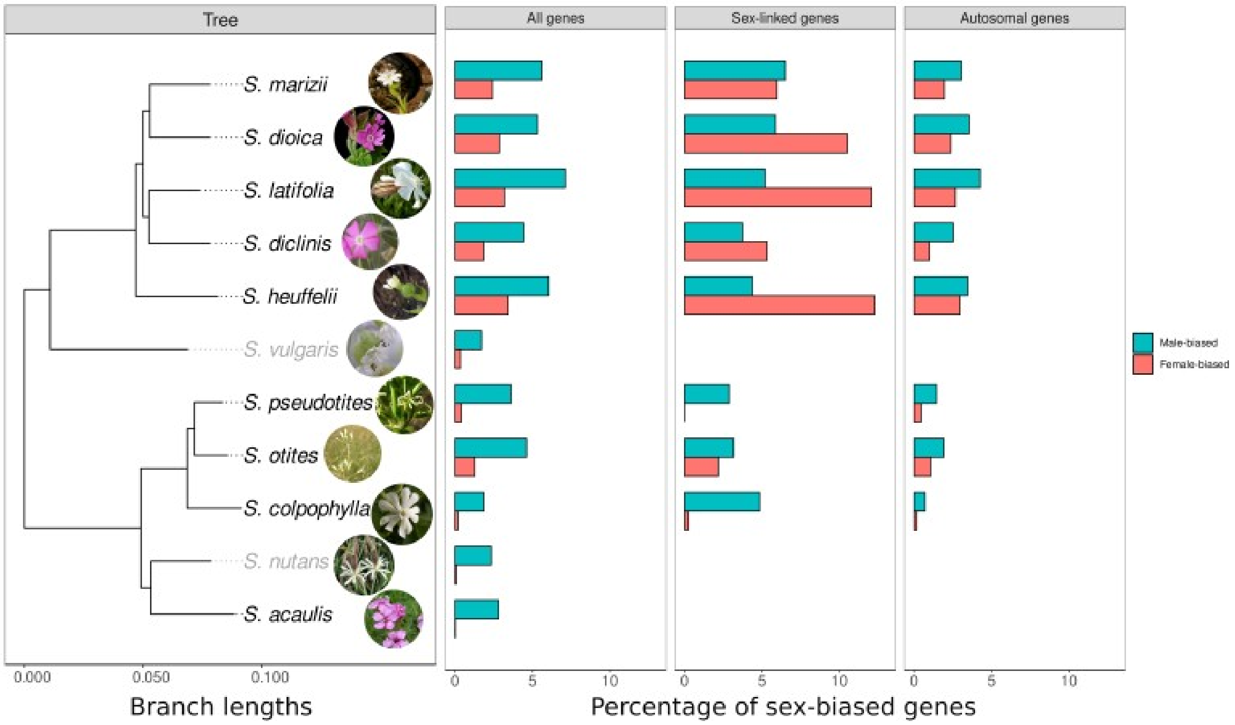

__**Supplementary Figure S2:**__ __Same as Figure 1 but with sex-biased genes inferred on a subset of the data with four males and four females for each species.__ Phylogenetic reconstruction (left panel) of the eleven *Silene* species used in this study and sex-biased gene proportions in each species (the outgroup *Dianthus chinensis* was removed for this figure, see Supplementary Figure S1 for a complete phylogeny). Gynodioecious and dioecious species names are written in grey and black, respectively. The proportion of expressed genes which are female-biased (red) or male-biased (blue) is shown for different gene subsets: all expressed genes (middle left panel), sex-linked genes (middle right panel, only for Otites and Melandrium sections) autosomal genes (right panel, only for Otites and Melandrium sections).

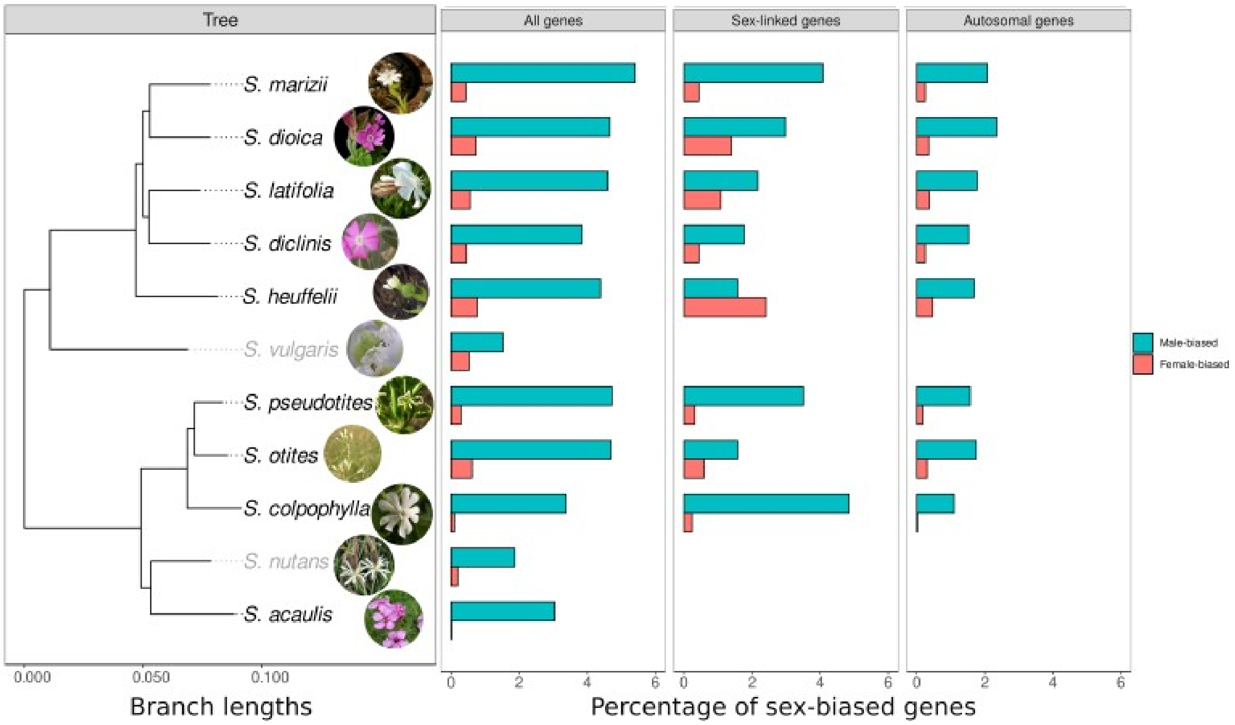

**<u>Supplementary Figure S3:</u>** <u>Same as Figure 1 but with a minimum foldchange of 4.</u> Phylogenetic reconstruction (left panel) of the eleven *Silene* species used in this study and sex-biased gene proportions in each species (the outgroup *Dianthus chinensis* was removed for this figure, see Supplementary Figure S1 for a complete phylogeny). Gynodioecious and dioecious species names are written in grey and black, respectively. The proportion of expressed genes which are female-biased (red) or male-biased (blue) is shown for different gene subsets: all expressed genes (middle left panel), sex-linked genes (middle right panel, only for Otites and Melandrium sections) autosomal genes (right panel, only for Otites and Melandrium sections).

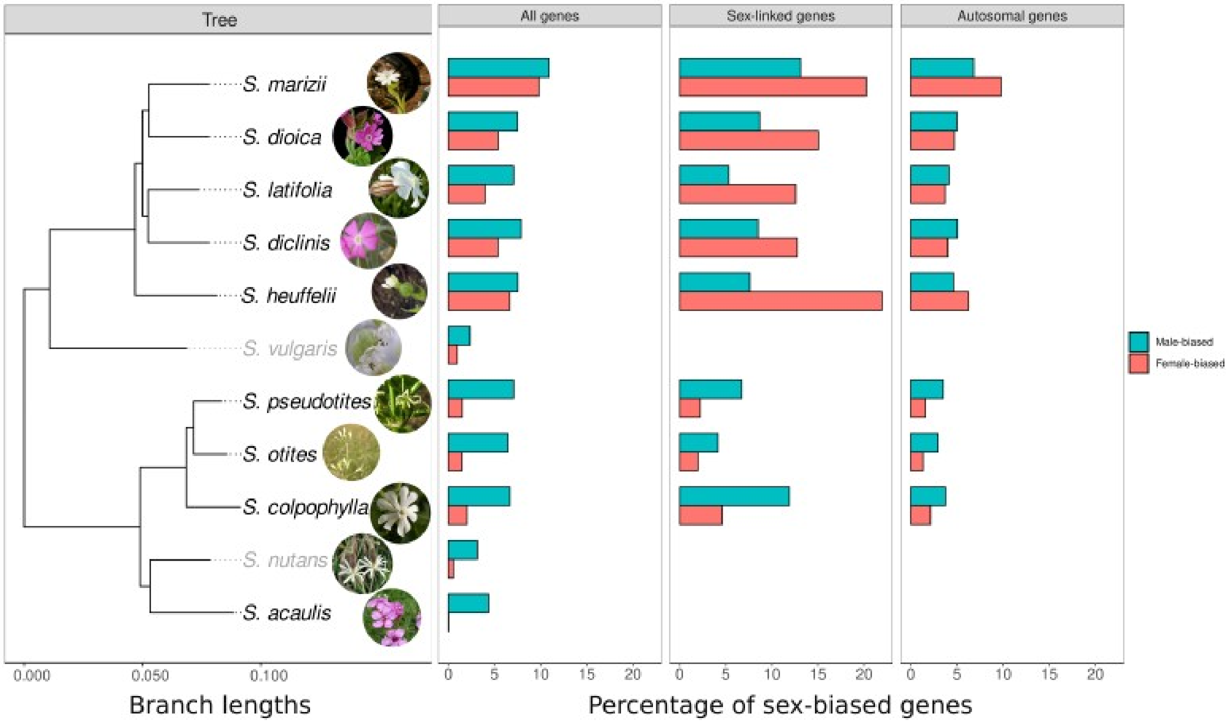

**Supplementary Figure S4:** Same as Figure 1 but with a maximal *p*-value=0.001. Phylogenetic reconstruction (left panel) of the eleven *Silene* species used in this study and sex-biased gene proportions in each species (the outgroup *Dianthus chinensis* was removed for this figure, see Supplementary Figure S1 for a complete phylogeny). Gynodioecious and dioecious species names are written in grey and black, respectively. The proportion of expressed genes which are female-biased (red) or male-biased (blue) is shown for different gene subsets: all expressed genes (middle left panel), sex-linked genes (middle right panel, only for Otites and Melandrium sections) autosomal genes (right panel, only for Otites and Melandrium sections).

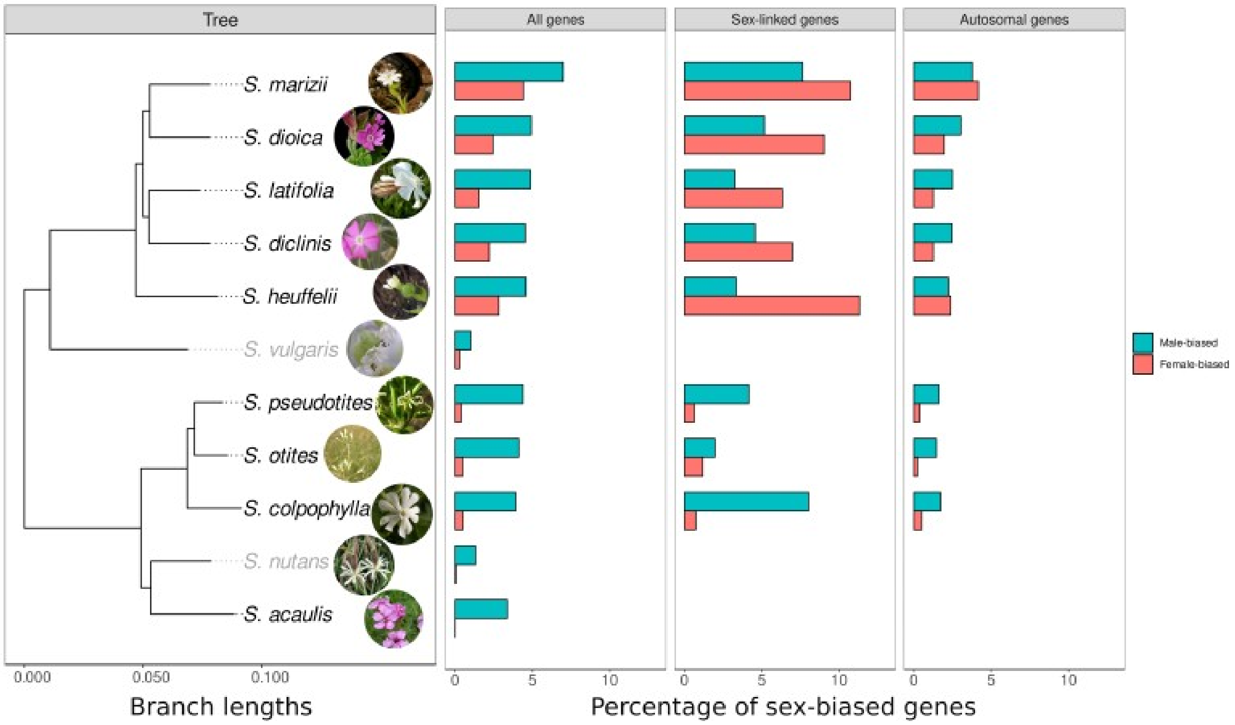

**<u>Supplementary Figure S5:</u>** <u>Same as Figure 1 but with a maximal *p*-value=0.00001.</u> Phylogenetic reconstruction (left panel) of the eleven *Silene* species used in this study and sex-biased gene proportions in each species (the outgroup *Dianthus chinensis* was removed for this figure, see Supplementary Figure S1 for a complete phylogeny). Gynodioecious and dioecious species names are written in grey and black, respectively. The proportion of expressed genes which are female-biased (red) or male-biased (blue) is shown for different gene subsets: all expressed genes (middle left panel), sex-linked genes (middle right panel, only for Otites and Melandrium sections) autosomal genes (right panel, only for Otites and Melandrium sections).

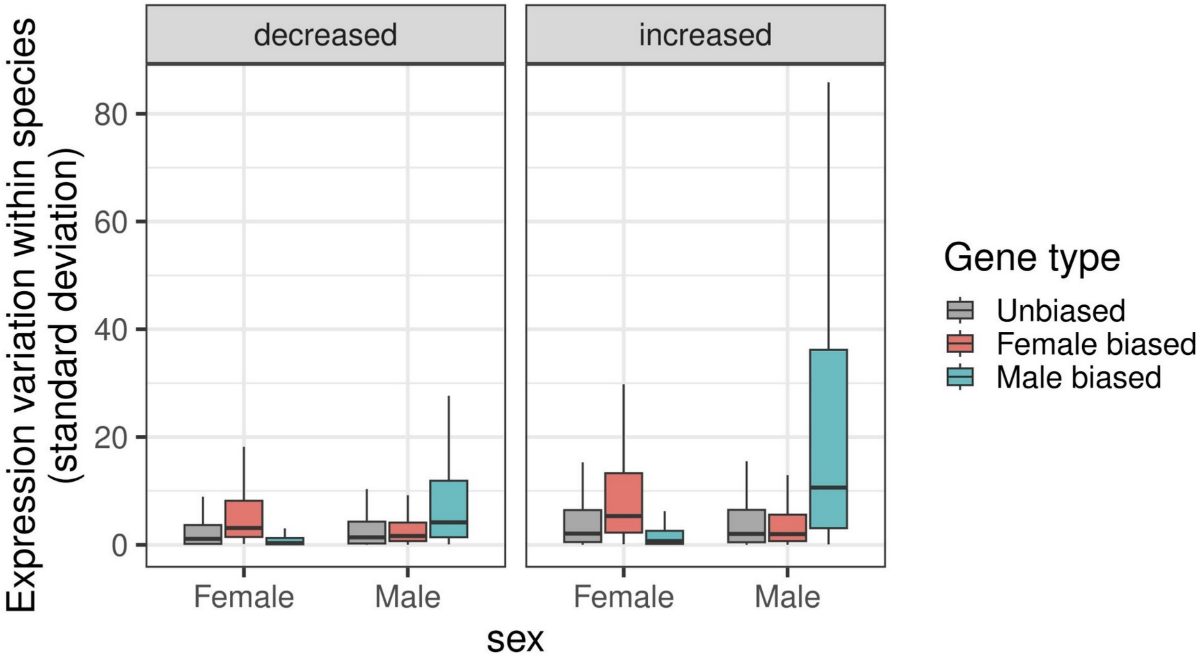

**Supplementary Figure S6:** Female and male expression variation within species for unbiased (gray), female-biased (red) and male-biased (blue) genes. The left panel shows genes whose expression decreased compared to the outgroup, while the right panel shows genes whose expression increased compared to the outgroup.

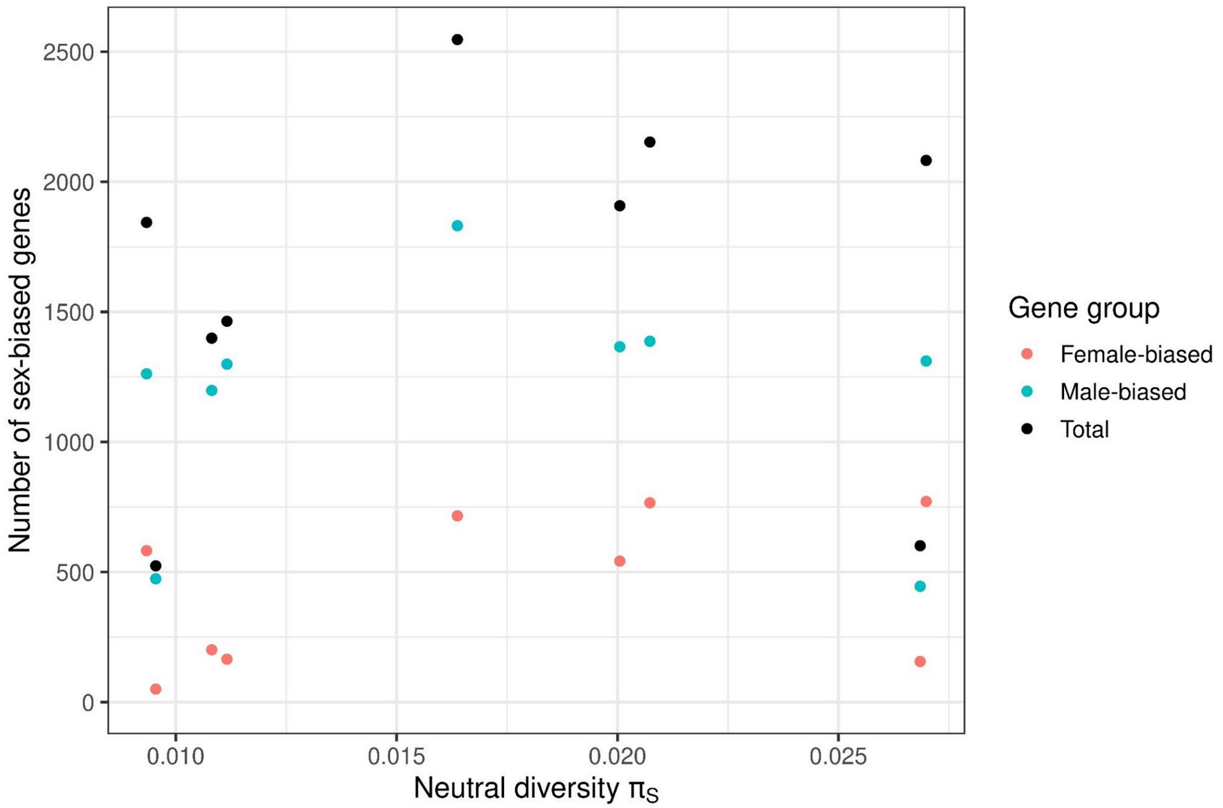

**Supplementary Figure S7:** Number of sex biased genes as a function of neutral diversity $\pi_S$ (used as a proxy for effective population size $N_e$). Total sex-biased gene numbers are shown in black, female-biased genes in red and male-biased genes in blue. None of the regressions were significant (Supplementary Table S6).

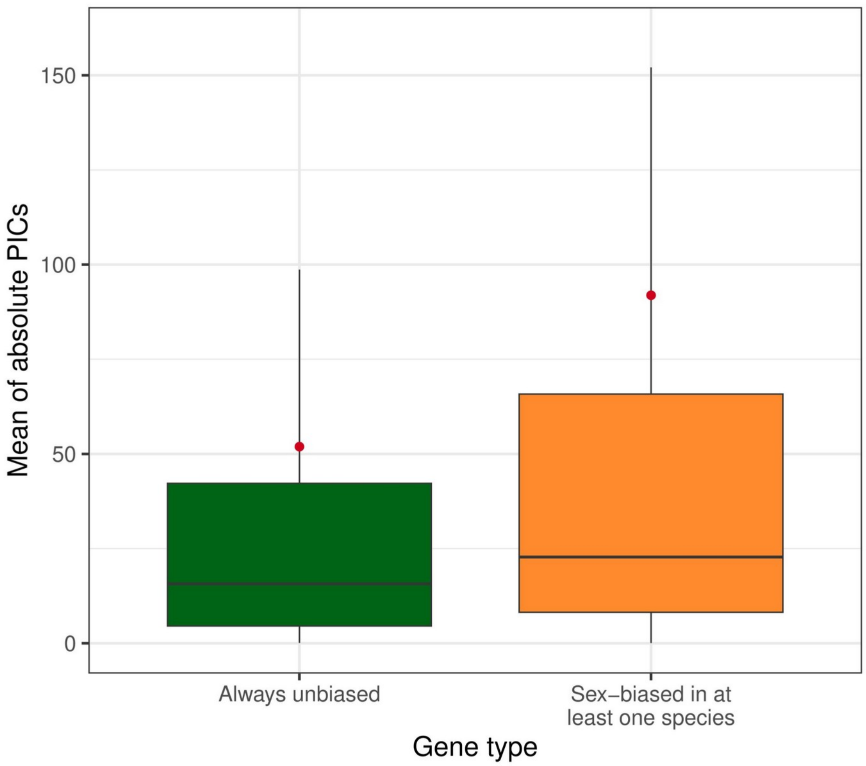

**<u>Supplementary Figure S8:</u>** Boxplot of mean absolute PIC values for gene expression change over time. Genes with at least one species showing SBGE are shown in orange and genes for which all species have unbiased expression in green. PICs for genes that are sex-biased in at least one species were computed after removing species for which the gene is sex-biased, in order to measure the gene expression change when it is not in a sex-biased state. The mean is indicated by the red dot.

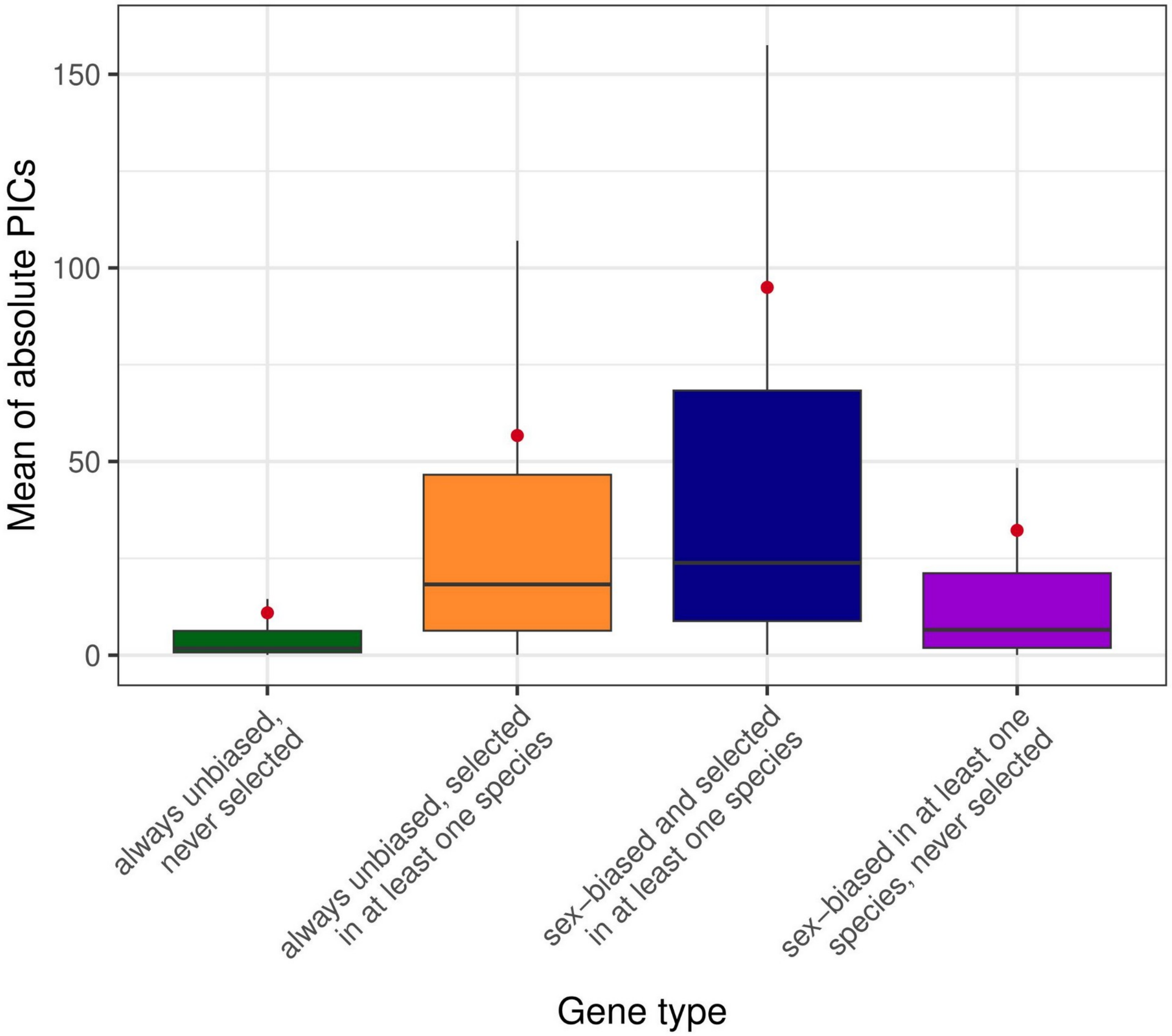

**Supplementary Figure S9:** Same as Supplementary Figure S7, but genes were further divided into showing positive selection ($\Delta_X$ higher than the quantile 75) in at least one species, or never being selected. PICs for genes that are sex-biased in at least one species were computed after removing species for which the gene is sex-biased, in order to measure the gene expression change when it is not in a sex-biased state. The mean is indicated by the red dot.

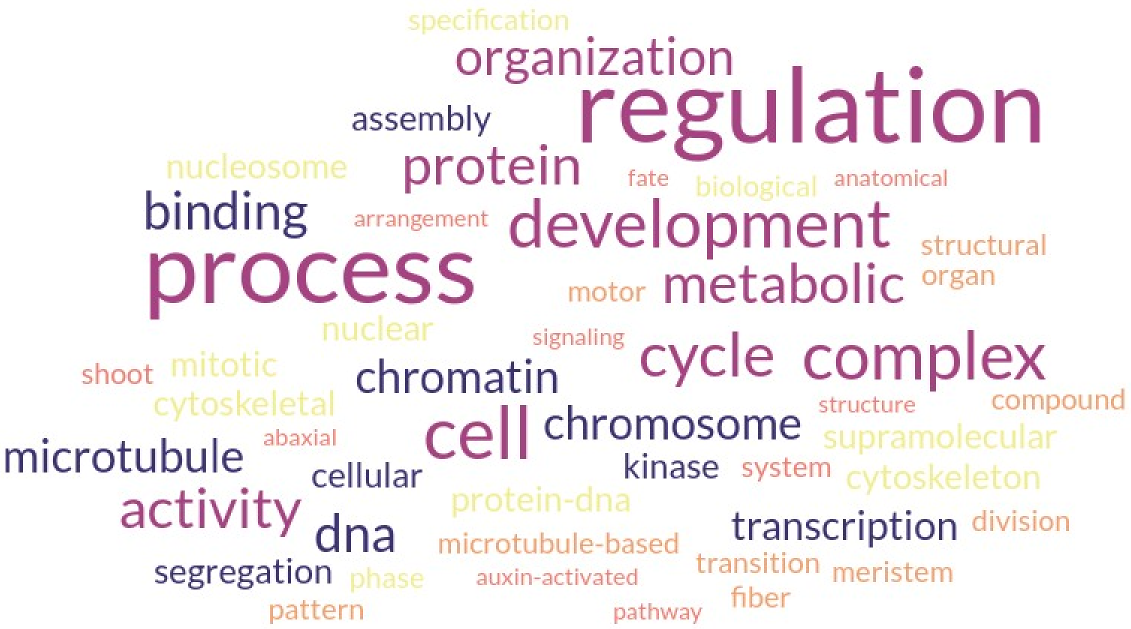

**Supplementary Figure S10:** Wordcloud of the functions enriched in female-biased genes while testing the enrichment against the whole annotation.

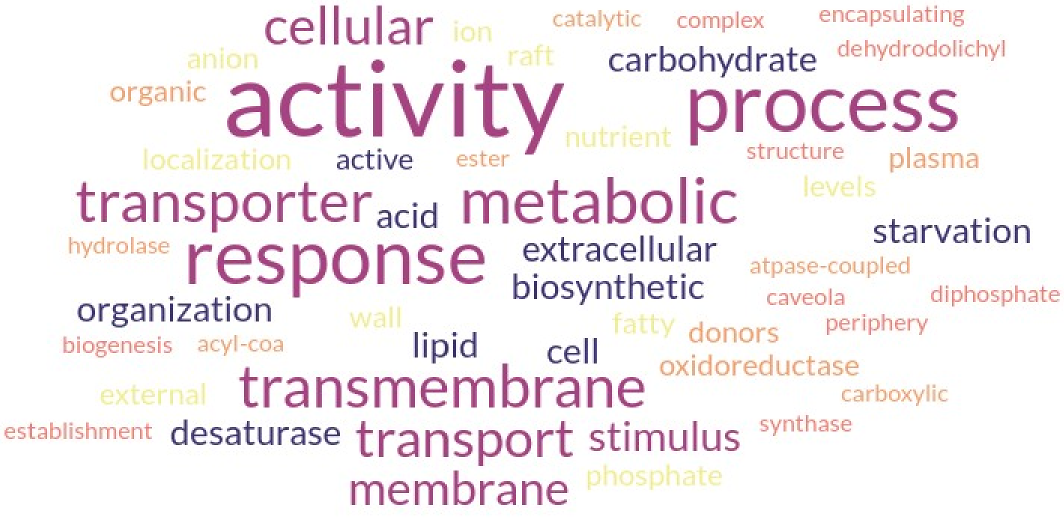

**Supplementary Figure S11:** Wordcloud of the functions enriched in male-biased genes while testing the enrichment against the whole annotation.

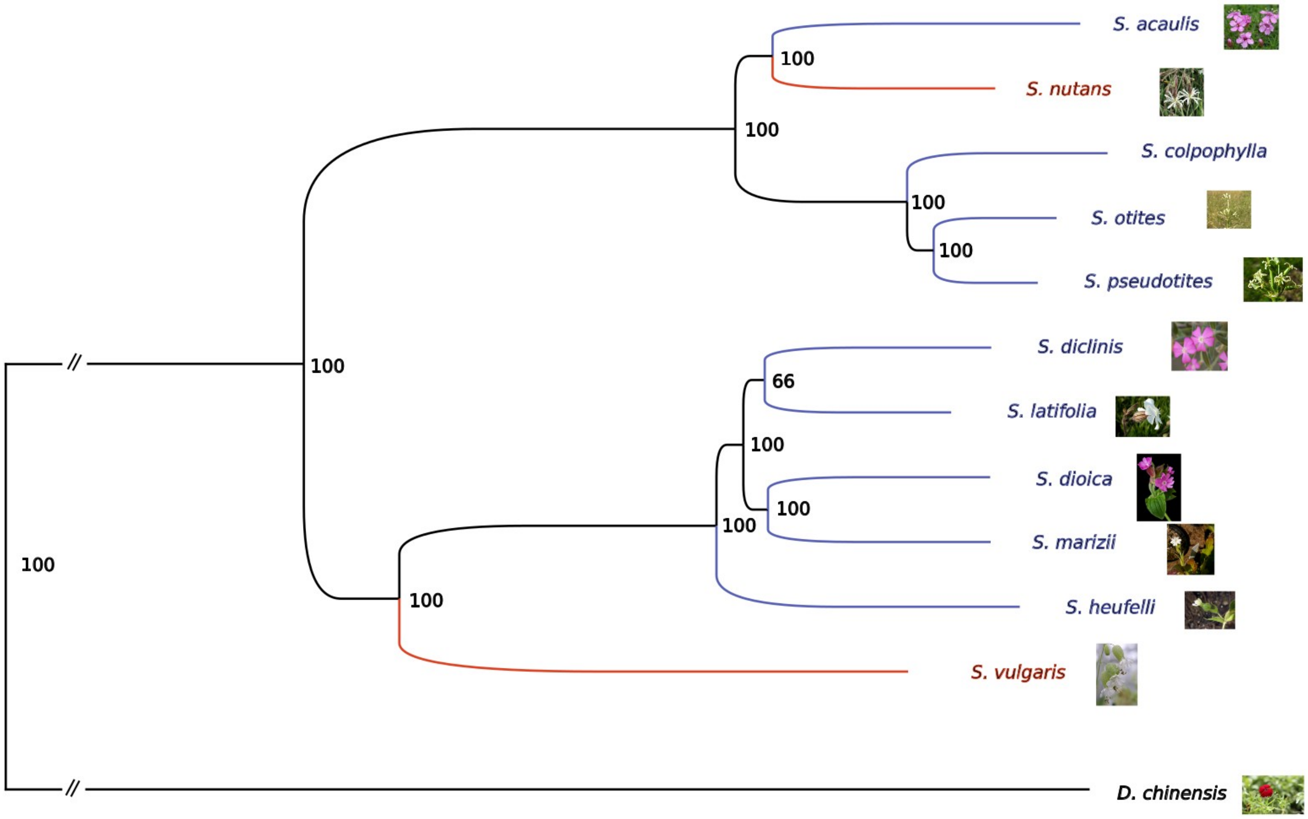

**Supplementary Figure S12:** Phylogenetic reconstruction of the 9 dioecious (in blue), the 2 gynodioecious species (in red) and *Dianthus chinensis* (in black, outgroup). The bootstrap values for 100 trials are indicated for each internal node.

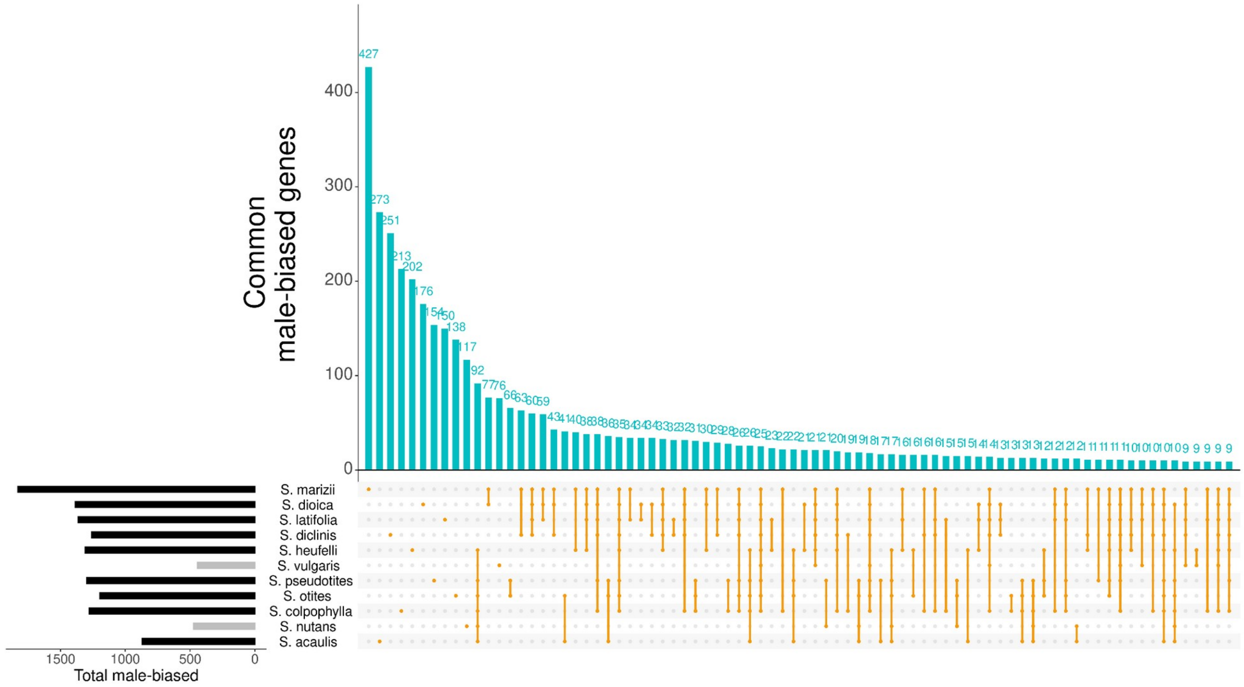

**Supplementary Figure S13:** Number of male-biased genes shared between different species, or unique to a single species. Only the 80 first interactions are shown here.

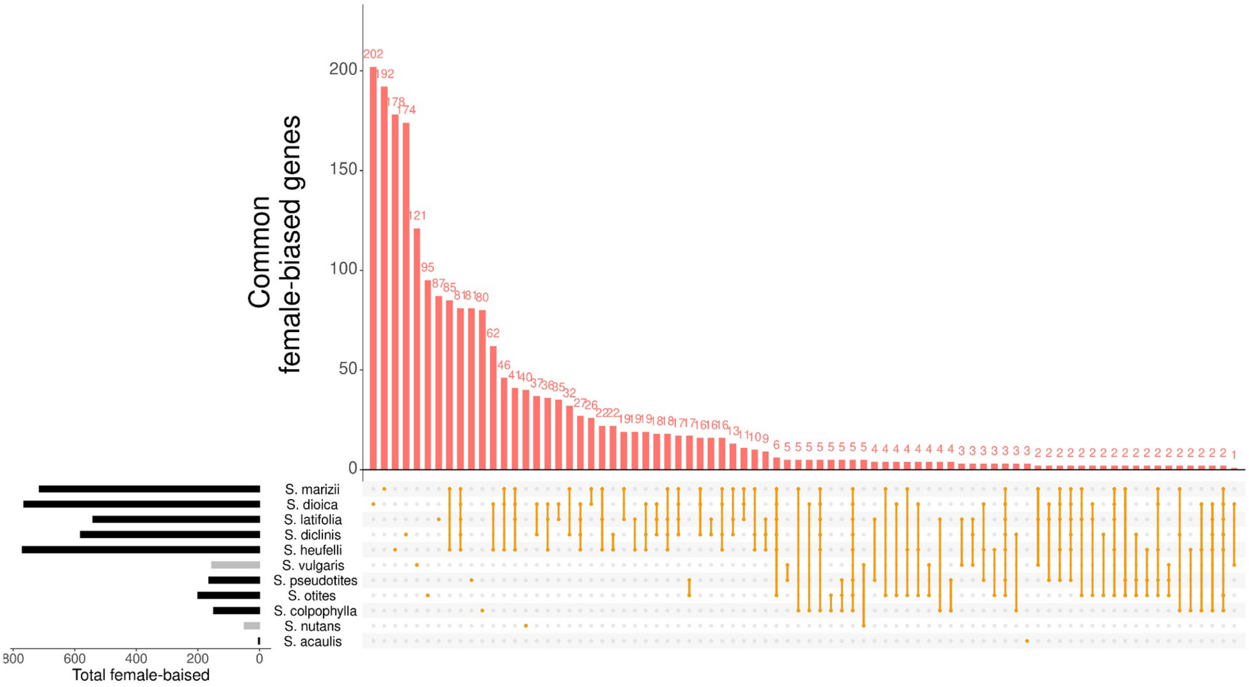

**Supplementary Figure S14:** Number of female-biased genes shared between different species, or unique to a single species. Only the 80 first interactions are shown here.

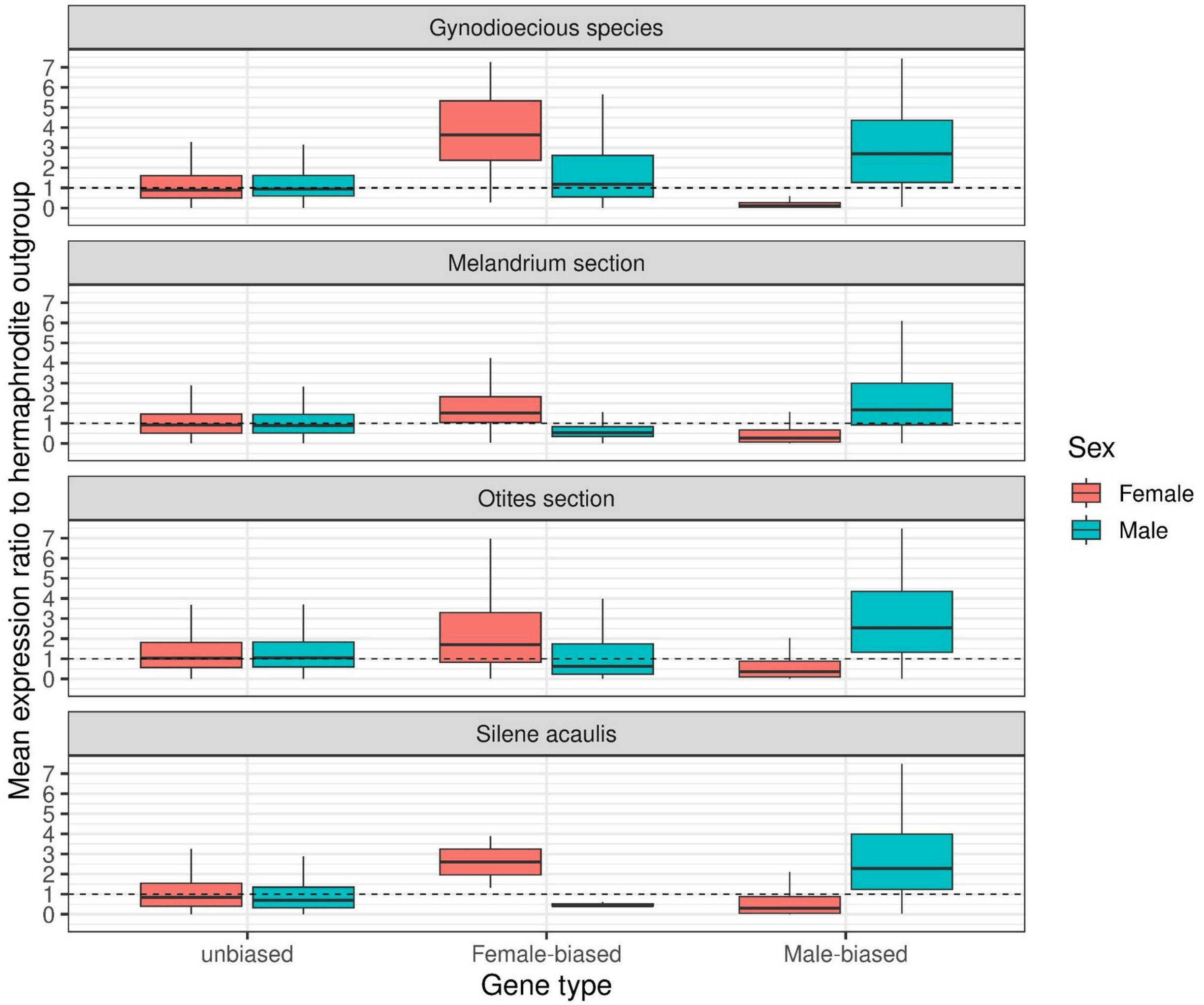

**Supplementary Figure S15:** Boxplot of the expression ratio between focal species and their hermaphrodite outgroup.

**<u>Supplementary Table S1:</u>** Statistics on de novo transcriptome assemblies for *S. vulgaris* and *S. nutans*. The transcriptomes were assembled using DRAP (version 1.92, Cabau et al. 2017). Busco (version 3.1.0; Simao et al, 2015) was run using eudicotyledons_odb10 (Creation date: 2017-12-01, number of species: 40, number of BUSCOs: 2121). The *S. vulgaris* transcriptome assembly is of higher quality.

| | ***S. vulgaris*** | ***S. nutans*** |
|---|---|---|
| **# contigs** | 31526 | 23836 |
| **N50** | 1494 | 1455 |
| **Score transrate** | 0.171 | 0.177 |
| **Complete busco** | 2010 (94.8 %) | 1706 (80.4 %) |
| **Complete and single** | 1798 (84.8 %) | 1592 (75.1 %) |
| **Complete and duplicated** | 212 (10.0 %) | 114 (5.4 %) |
| **Fragmented** | 35 (1.7 %) | 102 (4.8 %) |
| **Missing** | 76 (3.6 %) | 313 (1.5 %) |

**<u>Supplementary Table S2:</u>** Results of the mapping (in % of reads that were mapped properly-paired) on each *de novo* transcriptome assembly (one mapping iteration with gsnap). All hermaphrodite species have been mapped on *S. vulgaris* transcriptome only.

| **Species name** | **Tissue** | ***S. nutans* assembly** | | ***S. vulgaris* assembly** | |
|---|---|---|---|---|---|
| **Dioecious species** | | ♂ | ♀ | ♂ | ♀ |
| ***Silene acaulis*** | Flower buds | 60.5 | 58.3 | 48.9 | 49.6 |
| ***Silene colpophylla*** | Flower buds | 59.1 | 58.4 | 50.7 | 49.4 |
| ***Silene diclinis*** | Flower buds | 34.7 | 39.8 | 45.2 | 47.6 |
| ***Silene dioica*** | Flower buds | 41.1 | 43.6 | 49.3 | 51.6 |
| ***Silene heuffelii*** | Flower buds | 40.9 | 43.5 | 48.2 | 51.5 |
| ***Silene latifolia*** | Flower buds | 41.1 | 41.1 | 53.8 | 49.5 |
| ***Silene latifolia*** | Leaves | 55.0 | 56.3 | 46.6 | 48.1 |
| ***Silene marizii*** | Flower buds | 38.3 | 43.5 | 46.3 | 51.2 |
| ***Silene otites*** | Flower buds | 62.7 | 63.9 | 51.1 | 53.3 |
| ***Silene pseudotites*** | Flower buds | 62.3 | 64.3 | 51.08 | 53.3 |
| **Gynodioecious species** | | ⚥ | ♀ | ⚥ | ♀ |
| ***Silene nutans*** | Flower buds | 55.0 | 56.3 | 42.8 | 39.1 |
| ***Silene vulgaris*** | Flower buds | 42.3 | 41.4 | 55.6 | 53.5 |
| **Hermaphrodite species** | | ⚥ | | ⚥ | |
| ***Silene viscosa*** | Flower buds | NA | | 48.0 | |
| ***Silene viscosa*** | Leaves | NA | | 41.7 | |
| ***Silene paradoxa*** | Flower buds | NA | | 44.3 | |
| ***Dianthus chinensis*** | Seedlings | NA | | 9.0 | |

**<u>Supplementary Table S3:</u>** Number of expressed genes, male-biased genes, female-biased genes and total of sex-biased genes. For each of these categories we reported the number of autosomal ("Auto.") and sex-linked ("S-L") genes for the 8 dioecious species with a pair of sex chromosomes that have been identified.

| **Species** | **Number of expressed genes** | **Male-biased** (hermaphrodite-biased in gynodioecious species) | | | **Female-biased** | | | **Total sex-biased** | | |
|---|---|---|---|---|---|---|---|---|---|---|
| | | Total | Auto. | S-L | Total | Auto. | S-L | Total | Auto. | S-L |
| *Nanosilene* section | | | | | | | | | | |
| *S. acaulis* | 23541 | 872 | NA | NA | 4 | NA | NA | 876 | NA | NA |
| *Melandrium* section | | | | | | | | | | |
| *S. diclinis* | 23809 | 1262 | 346 | 73 | 582 | 190 | 104 | 1844 | 536 | 177 |
| *S. dioica* | 23552 | 1387 | 571 | 74 | 766 | 408 | 129 | 2153 | 979 | 203 |
| *S. heuffelii* | 23902 | 1311 | 469 | 44 | 771 | 458 | 125 | 2082 | 927 | 169 |
| *S. latifolia* | 24262 | 1366 | 438 | 42 | 542 | 269 | 98 | 1908 | 707 | 140 |
| *S. latifolia (leaves)* | 22851 | 27 | 4 | 7 | 9 | 1 | 3 | 36 | 5 | 10 |
| *S. marizii* | 23723 | 1831 | 530 | 84 | 716 | 307 | 75 | 2547 | 837 | 159 |
| *Otites* section | | | | | | | | | | |
| *S. colpophylla* | 27404 | 1281 | 240 | 34 | 150 | 56 | 2 | 1431 | 296 | 36 |
| *S. otites* | 24230 | 1198 | 191 | 13 | 201 | 53 | 7 | 1399 | 244 | 19 |
| *S. pseudotites* | 24070 | 1299 | 277 | 16 | 165 | 81 | 2 | 1464 | 358 | 18 |
| Gynodioecious species | | | | | | | | | | |
| *S. nutans* | 24608 | 474 | NA | NA | 50 | NA | NA | 524 | NA | NA |
| *S. vulgaris* | 29247 | 445 | NA | NA | 156 | NA | NA | 601 | NA | NA |

**<u>Supplementary Table S4:</u>** <u>Same Table as Supplementary Table S6 but with sex-biased genes inferred on a subset of the data with four males and four females for each species.</u> Generalized linear models (GLMs) between the number of sex-biased genes and the age of dioecy. Please refer to Materials and Methods (M&M) equations 2 and 4 for details. See Supplementary Figure S1 for a graphical representation. For the link between the number of sex-biased genes and the age of dioecy, no plateau was not necessary (i.e. no polynomial regression), unlike the analysis on the entire dataset presented in Figure 2 and Supplementary Table S5. The models correcting for phylogeny (M&M equation 4) are less flexible and do not allow to include an offset nor a negative binomial distribution and the polynomial model fails to converge.

| Number of sex-biased genes | Model type | Estimate | Standard Error | Z-value | p-value | $R^2$ |
|---|---|---|---|---|---|---|
| **Female-biased** | M&M equation 2 (GLM, negative binomial family, offset) | 0.21764 | 0.03219 | 6.761 | **1.37e-11** | **0.89** |
| **Total** | | 0.10530 | 0.01557 | 6.763 | **1.36e-11** | **0.84** |
| **Male-biased** | | 0.07887 | 0.01448 | 5.446 | **5.16e-08** | **0.75** |
| **Female-biased** | M&M Equation 4 (generalized least squares model correcting for phylogeny) | 1.291133 | 0.2621603 | 4.92498 | **8e-04** | NA |
| **Total** | | 0.523138 | 0.0838665 | 6.23774 | **2e-04** | NA |
| **Male-biased** | | 0.378137 | 0.0756771 | 4.99671 | **7e-04** | NA |

**<u>Supplementary Table S5:</u>** same as Supplementary Table S8, but on the subset of the data with four males and four females for every species.

| | | Expression increased compared to outgroup | | | | | | Expression decreased compared to outgroup | | | | | |
|---|---|---|---|---|---|---|---|---|---|---|---|---|---|
| | | Female ΔX | | | Male ΔX | | | Female ΔX | | | Male ΔX | | |
| Silene species | Sex-bias type | Proportion of sex-biased genes with high ΔX | Proportion of unbiased genes with high ΔX | corrected Chi2 p-value | Proportion of sex-biased genes with high ΔX | Proportion of unbiased genes with high ΔX | corrected Chi2 p-value | Proportion of sex-biased genes with high ΔX | Proportion of unbiased genes with high ΔX | corrected Chi2 p-value | Proportion of sex-biased genes with high ΔX | Proportion of unbiased genes with high ΔX | corrected Chi2 p-value |
| ***acaulis*** | **Female-biased genes** | 0,17 | 0,06 | 8,69E-01 | 0,00 | 0,06 | 1,00E+00 | 0,67 | 0,40 | 8,42E-01 | 0,67 | 0,39 | 4,61E-01 |
| ***colpophylla*** | | 0,30 | 0,20 | 2,34E-01 | 0,06 | 0,13 | 4,93E-01 | 0,60 | 0,45 | 9,07E-01 | 0,45 | 0,38 | 8,42E-01 |
| ***diclinis*** | | **0,24** | **0,13** | **8,85E-08** | 0,10 | 0,10 | 1,00E+00 | **0,25** | **0,38** | **1,65E-02** | **0,59** | **0,36** | **4,38E-16** |
| ***dioica*** | | **0,32** | **0,15** | **7,47E-26** | 0,11 | 0,09 | 4,98E-01 | **0,20** | **0,37** | **7,37E-04** | **0,59** | **0,34** | **1,40E-30** |
| ***heuffelli*** | | **0,18** | **0,15** | **4,02E-02** | 0,14 | 0,09 | 1,76E-01 | **0,26** | **0,39** | **6,55E-03** | **0,48** | **0,36** | **3,32E-10** |
| ***latifolia*** | | **0,31** | **0,15** | **1,62E-25** | 0,09 | 0,11 | 5,60E-01 | **0,23** | **0,36** | **3,02E-03** | **0,50** | **0,34** | **5,18E-16** |
| ***marizii*** | | **0,18** | **0,12** | **5,58E-04** | 0,09 | 0,12 | 5,23E-01 | **0,20** | **0,37** | **3,27E-04** | **0,60** | **0,37** | **2,69E-21** |
| ***nutans*** | | **0,23** | **0,07** | **1,91E-02** | 0,06 | 0,05 | 1,00E+00 | 0,00 | 0,46 | 4,15E-01 | 0,67 | 0,33 | 1,06E-01 |
| ***otites*** | | 0,14 | 0,14 | 1,00E+00 | 0,06 | 0,11 | 3,99E-01 | 0,40 | 0,39 | 1,00E+00 | **0,68** | **0,38** | **7,87E-18** |
| ***pseudotites*** | | 0,10 | 0,13 | 8,14E-01 | 0,13 | 0,09 | 8,42E-01 | 0,60 | 0,41 | 8,21E-02 | **0,63** | **0,37** | **4,76E-05** |
| ***vulgaris*** | | 0,17 | 0,15 | 8,42E-01 | **0,01** | **0,13** | **1,65E-02** | 0,00 | 0,39 | 8,25E-01 | 0,34 | 0,45 | 3,65E-01 |
| ***acaulis*** | **Male-biased genes** | 0,04 | 0,06 | 8,42E-01 | **0,12** | **0,06** | **3,09E-09** | **0,68** | **0,40** | **5,77E-33** | **0,24** | **0,39** | **2,82E-02** |
| ***colpophylla*** | | **0,13** | **0,20** | **9,46E-03** | **0,03** | **0,13** | **1,07E-09** | **0,58** | **0,45** | **1,86E-04** | 0,26 | 0,38 | 1,76E-01 |
| ***diclinis*** | | 0,16 | 0,13 | 6,09E-01 | **0,04** | **0,10** | **1,44E-07** | **0,74** | **0,38** | **8,33E-100** | **0,21** | **0,36** | **2,92E-06** |
| ***dioica*** | | **0,09** | **0,15** | **2,52E-02** | 0,08 | 0,09 | 8,69E-01 | **0,72** | **0,37** | **2,21E-99** | 0,34 | 0,34 | 1,00E+00 |
| ***heuffelli*** | | **0,06** | **0,15** | **3,13E-05** | **0,04** | **0,09** | **1,88E-08** | **0,64** | **0,39** | **2,01E-57** | 0,35 | 0,36 | 9,01E-01 |
| ***latifolia*** | | **0,04** | **0,15** | **5,49E-06** | **0,07** | **0,11** | **2,10E-06** | **0,71** | **0,36** | **2,22E-147** | **0,23** | **0,34** | **8,67E-06** |

| | | | | | | | | | | | | | |
|---|---|---|---|---|---|---|---|---|---|---|---|---|---|
| ***marizii*** | | 0,08 | 0,12 | 1,47E-01 | **0,09** | **0,12** | **1,96E-02** | **0,69** | **0,37** | **2,70E-92** | **0,19** | **0,37** | **7,03E-09** |
| ***nutans*** | | 0,11 | 0,07 | 9,07E-01 | **0,13** | **0,05** | **1,36E-17** | **0,88** | **0,46** | **1,59E-77** | **0,04** | **0,33** | **2,11E-04** |
| ***otites*** | | **0,06** | **0,14** | **3,02E-03** | **0,03** | **0,11** | **3,19E-13** | **0,69** | **0,39** | **1,75E-61** | **0,26** | **0,38** | **4,02E-02** |
| ***pseudotites*** | | **0,05** | **0,13** | **1,01E-02** | **0,03** | **0,09** | **7,91E-09** | **0,68** | **0,41** | **1,61E-42** | **0,12** | **0,37** | **2,64E-03** |
| ***vulgaris*** | | **0,04** | **0,15** | **4,37E-03** | **0,07** | **0,13** | **7,13E-04** | **0,78** | **0,39** | **9,11E-49** | **0,14** | **0,45** | **2,80E-03** |

**Supplementary Table S6:** Generalized linear models (GLMs) between the number of sex-biased genes and the age of dioecy. Please refer to Materials and Methods (M&M) equations 1, 2 and 4 for details. See Figure 2 for a graphical representation. For polynomial regressions of degree 2 ($y=a+bx+cx^2$, equation 1), two lines are indicated in the table, the first line for the effect of x and the second line for the effect of $x^2$, where x stands for the age of dioecy. The effect of x is significantly positive, showing a positive correlation between the number of sex-biased genes and the age of dioecy. The effect of $x^2$ is negative, allowing for a plateau in the correlation (Figure 2). For female-biased genes, a plateau was not necessary. The models correcting for phylogeny (M&M equation 4) are less flexible and do not allow to include an offset nor a negative binomial distribution and the polynomial model fails to converge.

| Number of sex-biased genes | Model type | Estimate | Standard Error | Z-value | p-value | $R^2$ |
|---|---|---|---|---|---|---|
| **Female-biased** | M&M equation 2 (GLM, negative binomial family, offset) | 0.20245 | 0.02808 | 7.209 | **5.62e-13** | **0.90** |
| **Total** | M&M equation 1 (polynomial GLM, negative binomial family, offset) | x: 0.48515 | 0.073798 | 6.57 | **4.90e-11** | **0.96** |
| | | $x^2$: -0.03280 | 0.005891 | -5.57 | **2.58e-08** | |
| **Male-biased** | | x: 0.50904 | 0.082369 | 6.180 | **6.41e-10** | **0.92** |
| | | $x^2$: -0.03693 | 0.006576 | -5.615 | **1.96e-08** | |
| **Female-biased** | M&M Equation 4 (generalized least squares model correcting for phylogeny) | 1.204517 | 0.344717 | 3.49422 | **0.0068** | NA |
| **Total** | | 0.526127 | 0.115804 | 4.54326 | **0.0014** | NA |
| **Male-biased** | | 0.440711 | 0.144519 | 3.04949 | **0.0138** | NA |

**Supplementary Table S7:** Comparison of the proportion of sex-biased and unbiased genes under selection. Genes were defined as under selection if their $\Delta_X$ values were higher than the 75 quantile. A Chi-square test was used to test for the enrichment of sex-biased genes in positive selection compared to unbiased genes. Text in black bold indicates a significant enrichment of sex biased genes in selection compared to unbiased genes. Text in red shows a significant depletion of sex biased genes in selection compared to unbiased genes.

| Silene species | Sex-bias type | Female ΔX | | | Male ΔX | | |
|---|---|---|---|---|---|---|---|
| | | Proportion of sex-biased genes with high ΔX | Proportion of unbiased genes with high ΔX | corrected Chi2 p-value | Proportion of sex-biased genes with high ΔX | Proportion of unbiased genes with high ΔX | corrected Chi2 p-value |
| ***acaulis*** | Female-biased genes | 0,00 | 0,23 | 6,21E-01 | 0,25 | 0,26 | 1,00E+00 |
| ***colpophylla*** | | 0,34 | 0,31 | 4,88E-01 | 0,23 | 0,19 | 3,71E-01 |
| ***diclinis*** | | **0,17** | **0,25** | **1,61E-05** | **0,46** | **0,24** | **7,18E-34** |
| ***dioica*** | | 0,26 | 0,27 | 4,33E-01 | **0,45** | **0,21** | **2,59E-55** |
| ***heuffelli*** | | **0,20** | **0,27** | **1,61E-05** | **0,44** | **0,22** | **4,14E-46** |
| ***latifolia*** | | 0,21 | 0,19 | 2,29E-01 | **0,56** | **0,29** | **1,52E-41** |
| ***marizii*** | | **0,20** | **0,25** | **2,42E-03** | **0,49** | **0,23** | **1,35E-57** |
| ***nutans*** | | **0,10** | **0,33** | **1,24E-03** | 0,12 | 0,16 | 5,42E-01 |
| ***otites*** | | 0,22 | 0,25 | 4,49E-01 | **0,52** | **0,24** | **1,10E-19** |
| ***pseudotites*** | | **0,21** | **0,28** | **5,79E-02** | **0,43** | **0,22** | **5,59E-11** |
| ***vulgaris*** | | **0,16** | **0,27** | **4,40E-03** | **0,13** | **0,23** | **7,78E-03** |
| ***acaulis*** | Male-biased genes | **0,52** | **0,23** | **9,30E-83** | **0,05** | **0,26** | **1,44E-46** |
| ***colpophylla*** | | **0,44** | **0,31** | **3,63E-23** | **0,06** | **0,19** | **1,64E-32** |
| ***diclinis*** | | **0,62** | **0,25** | **4,14E-185** | **0,08** | **0,24** | **3,57E-36** |
| ***dioica*** | | **0,61** | **0,27** | **7,96E-162** | **0,08** | **0,21** | **6,57E-33** |
| ***heuffelli*** | | **0,49** | **0,27** | **8,18E-67** | **0,06** | **0,22** | **6,47E-43** |
| ***latifolia*** | | **0,67** | **0,19** | **0,00E+00** | **0,19** | **0,29** | **3,92E-15** |
| ***marizii*** | | **0,58** | **0,25** | **4,29E-207** | **0,07** | **0,23** | **1,11E-58** |
| ***nutans*** | | **0,95** | **0,33** | **4,14E-171** | **0,03** | **0,16** | **8,64E-15** |
| ***otites*** | | **0,58** | **0,25** | **3,07E-139** | **0,03** | **0,24** | **1,92E-62** |
| ***pseudotites*** | | **0,58** | **0,28** | **4,30E-118** | **0,03** | **0,22** | **2,93E-55** |
| ***vulgaris*** | | **0,65** | **0,27** | **1,12E-69** | **0,09** | **0,23** | **2,60E-12** |

**Supplementary Table S8:** same as Supplementary Table S7 after splitting genes into increased (left columns) or decreased (right columns) expression compared to the hermaphrodite outgroup.

| | | Expression increased compared to outgroup | | | | | | Expression decreased compared to outgroup | | | | | |
|---|---|---|---|---|---|---|---|---|---|---|---|---|---|
| | | Female ΔX | | | Male ΔX | | | Female ΔX | | | Male ΔX | | |
| **Silene species** | **Sex-bias type** | **Proportion of sex-biased genes with high ΔX** | **Proportion of unbiased genes with high ΔX** | **corrected Chi2 p-value** | **Proportion of sex-biased genes with high ΔX** | **Proportion of unbiased genes with high ΔX** | **corrected Chi2 p-value** | **Proportion of sex-biased genes with high ΔX** | **Proportion of unbiased genes with high ΔX** | **corrected Chi2 p-value** | **Proportion of sex-biased genes with high ΔX** | **Proportion of unbiased genes with high ΔX** | **corrected Chi2 p-value** |
| ***acaulis*** | **Female-biased genes** | 0,00 | 0,04 | 1,00E+00 | 0,00 | 0,03 | 1,00E+00 | 0,00 | 0,41 | NA | 0,33 | 0,42 | 1,00E+00 |
| ***colpophylla*** | | **0,31** | **0,23** | **4,21E-02** | 0,11 | 0,11 | 1,00E+00 | 0,60 | 0,47 | 6,12E-01 | **0,58** | **0,36** | **1,34E-02** |
| ***diclinis*** | | 0,14 | 0,13 | 3,73E-01 | 0,08 | 0,10 | 6,79E-01 | **0,24** | **0,38** | **1,15E-03** | **0,57** | **0,38** | **4,40E-15** |
| ***dioica*** | | **0,25** | **0,14** | **7,11E-14** | 0,07 | 0,06 | 1,00E+00 | **0,27** | **0,39** | **4,97E-03** | **0,55** | **0,35** | **2,07E-24** |
| ***heuffelli*** | | **0,19** | **0,15** | **1,31E-02** | 0,10 | 0,08 | 4,94E-01 | **0,28** | **0,40** | **1,26E-02** | **0,53** | **0,36** | **6,11E-16** |
| ***latifolia*** | | **0,21** | **0,06** | **1,99E-32** | 0,19 | 0,16 | 6,12E-01 | **0,20** | **0,32** | **2,39E-02** | **0,66** | **0,42** | **8,02E-22** |
| ***marizii*** | | **0,19** | **0,12** | **2,23E-05** | 0,11 | 0,09 | 8,51E-01 | **0,24** | **0,38** | **4,12E-04** | **0,60** | **0,38** | **1,87E-25** |
| ***nutans*** | | 0,10 | 0,09 | 1,00E+00 | 0,00 | 0,03 | 7,34E-01 | 0,00 | 0,51 | 1,00E+00 | 0,35 | 0,29 | 8,81E-01 |
| ***otites*** | | 0,13 | 0,12 | 8,51E-01 | 0,06 | 0,09 | 6,79E-01 | 0,51 | 0,40 | 1,93E-01 | **0,69** | **0,41** | **9,55E-12** |
| ***pseudotites*** | | 0,14 | 0,12 | 8,52E-01 | 0,06 | 0,07 | 1,00E+00 | 0,40 | 0,42 | 1,00E+00 | **0,61** | **0,37** | **9,79E-07** |
| ***vulgaris*** | | 0,16 | 0,18 | 8,51E-01 | 0,02 | 0,11 | 1,06E-02 | 0,00 | 0,42 | 7,77E-01 | 0,35 | 0,42 | 4,86E-01 |
| ***acaulis*** | **Male-biased genes** | 0,04 | 0,04 | 1,00E+00 | 0,02 | 0,03 | 1,71E-01 | **0,65** | **0,41** | **7,72E-36** | **0,21** | **0,42** | **6,48E-05** |
| ***colpophylla*** | | **0,11** | **0,23** | **6,04E-10** | **0,01** | **0,11** | **1,48E-22** | **0,66** | **0,47** | **2,20E-22** | 0,29 | 0,36 | 7,52E-02 |
| ***diclinis*** | | 0,07 | 0,13 | 5,13E-02 | **0,04** | **0,10** | **2,18E-08** | **0,73** | **0,38** | **1,06E-108** | **0,22** | **0,38** | **7,48E-08** |
| ***dioica*** | | **0,05** | **0,14** | **4,46E-04** | **0,03** | **0,06** | **2,63E-05** | **0,74** | **0,39** | **2,56E-107** | 0,29 | 0,35 | 1,29E-01 |
| ***heuffelli*** | | **0,04** | **0,15** | **2,51E-07** | **0,02** | **0,08** | **1,38E-11** | **0,64** | **0,40** | **4,74E-48** | **0,26** | **0,36** | **6,88E-03** |
| ***latifolia*** | | **0,02** | **0,06** | **3,61E-02** | 0,15 | 0,16 | 9,15E-01 | **0,77** | **0,32** | **1,62E-207** | **0,31** | **0,42** | **2,87E-04** |
| ***marizii*** | | **0,07** | **0,12** | **3,52E-03** | **0,04** | **0,09** | **3,88E-12** | **0,71** | **0,38** | **4,57E-125** | **0,19** | **0,38** | **5,55E-14** |
| ***nutans*** | | 0,00 | 0,09 | 1,00E+00 | **0,01** | **0,03** | **7,78E-03** | **0,96** | **0,51** | **1,91E-80** | **0,14** | **0,29** | **9,76E-03** |
| ***otites*** | | **0,04** | **0,12** | **1,87E-03** | **0,02** | **0,09** | **3,01E-15** | **0,71** | **0,40** | **7,84E-80** | **0,25** | **0,41** | **2,70E-02** |
| ***pseudotites*** | | **0,05** | **0,12** | **1,33E-03** | **0,01** | **0,07** | **5,28E-13** | **0,69** | **0,42** | **1,76E-64** | **0,24** | **0,37** | **6,88E-03** |
| ***vulgaris*** | | **0,04** | **0,18** | **3,82E-03** | 0,08 | 0,11 | 1,51E-01 | **0,80** | **0,42** | **4,94E-44** | **0,13** | **0,42** | **6,25E-03** |

**Supplementary Table S9:** same as Supplementary Table S8, but considering genes as under positive selection when ΔX is higher than 10 (as in Scharman *et al.* 2021) instead of higher than the species 75 quantile (as in Zemp *et al.* 2016).

| Silene species | Sex-bias type | Expression increased compared to outgroup | | | | | | Expression decreased compared to outgroup | | | | | |
|---|---|---|---|---|---|---|---|---|---|---|---|---|---|
| | | Female ΔX | | | Male ΔX | | | Female ΔX | | | Male ΔX | | |
| | | Proportion of sex-biased genes with high ΔX | Proportion of unbiased genes with high ΔX | corrected Chi2 p-value | Proportion of sex-biased genes with high ΔX | Proportion of unbiased genes with high ΔX | corrected Chi2 p-value | Proportion of sex-biased genes with high ΔX | Proportion of unbiased genes with high ΔX | corrected Chi2 p-value | Proportion of sex-biased genes with high ΔX | Proportion of unbiased genes with high ΔX | corrected Chi2 p-value |
| ***acaulis*** | | 0,00 | 0,00 | 1,00E+00 | 0,00 | 0,00 | 1,00E+00 | NA | 0,15 | NA | 0,00 | 0,15 | 1,00E+00 |
| ***colpophylla*** | | 0,04 | 0,02 | 4,18E-01 | 0,00 | 0,01 | 1,00E+00 | 0,13 | 0,25 | 7,41E-01 | **0,39** | **0,16** | **1,23E-03** |
| ***diclinis*** | | 0,01 | 0,01 | 1,00E+00 | 0,02 | 0,01 | 8,79E-01 | **0,10** | **0,18** | **4,83E-02** | **0,29** | **0,17** | **6,62E-10** |
| ***dioica*** | | 0,01 | 0,01 | 8,63E-01 | 0,00 | 0,00 | 1,00E+00 | **0,07** | **0,16** | **1,29E-02** | **0,20** | **0,14** | **1,45E-04** |
| ***heuffelli*** | | 0,01 | 0,01 | 5,66E-01 | 0,01 | 0,00 | 1,00E+00 | 0,11 | 0,18 | 2,00E-01 | **0,19** | **0,15** | **4,03E-02** |
| ***latifolia*** | **Female-biased genes** | **0,02** | **0,01** | **3,23E-02** | 0,00 | 0,02 | 5,29E-01 | 0,07 | 0,14 | 1,04E-01 | **0,31** | **0,18** | **6,53E-12** |
| ***marizii*** | | 0,01 | 0,01 | 1,00E+00 | 0,01 | 0,01 | 1,00E+00 | **0,11** | **0,20** | **1,72E-02** | **0,34** | **0,18** | **1,28E-19** |
| ***nutans*** | | 0,00 | 0,00 | 1,00E+00 | 0,00 | 0,00 | | 0,00 | 0,16 | 1,00E+00 | 0,06 | 0,05 | 1,00E+00 |
| ***otites*** | | 0,00 | 0,00 | 1,00E+00 | 0,00 | 0,00 | 1,00E+00 | 0,18 | 0,13 | 7,05E-01 | **0,37** | **0,13** | **6,31E-16** |
| ***pseudotites*** | | 0,00 | 0,00 | 1,00E+00 | 0,00 | 0,00 | 1,00E+00 | 0,17 | 0,16 | 1,00E+00 | **0,30** | **0,13** | **2,14E-06** |
| ***vulgaris*** | | 0,00 | 0,01 | 8,79E-01 | 0,00 | 0,00 | 1,00E+00 | 0,00 | 0,10 | 1,00E+00 | 0,13 | 0,07 | 3,50E-01 |
| ***acaulis*** | | 0,00 | 0,00 | 1,00E+00 | 0,00 | 0,00 | 1,00E+00 | **0,44** | **0,15** | **7,98E-83** | 0,07 | 0,15 | 6,67E-02 |
| ***colpophylla*** | | 0,01 | 0,02 | 2,87E-01 | 0,00 | 0,01 | 7,53E-02 | **0,50** | **0,25** | **6,85E-51** | 0,17 | 0,16 | 1,00E+00 |
| ***diclinis*** | | 0,00 | 0,01 | 1,00E+00 | 0,00 | 0,01 | 2,35E-01 | **0,52** | **0,18** | **4,57E-157** | **0,11** | **0,17** | **4,03E-02** |
| ***dioica*** | | 0,00 | 0,01 | 6,86E-01 | 0,00 | 0,00 | 5,99E-01 | **0,49** | **0,16** | **2,14E-154** | 0,11 | 0,14 | 6,24E-01 |
| ***heuffelli*** | | 0,00 | 0,01 | 5,29E-01 | 0,00 | 0,00 | 5,29E-01 | **0,43** | **0,18** | **9,46E-82** | 0,09 | 0,15 | 7,85E-02 |
| ***latifolia*** | **Male-biased genes** | 0,00 | 0,01 | 8,79E-01 | **0,00** | **0,02** | **3,04E-03** | **0,56** | **0,14** | **6,80E-280** | 0,13 | 0,18 | 9,21E-02 |
| ***marizii*** | | 0,01 | 0,01 | 5,29E-01 | **0,00** | **0,01** | **6,32E-03** | **0,52** | **0,20** | **3,78E-161** | **0,08** | **0,18** | **1,15E-06** |
| ***nutans*** | | 0,00 | 0,00 | 1,00E+00 | 0,00 | 0,00 | NA | **0,74** | **0,16** | **6,05E-234** | 0,01 | 0,05 | 4,18E-01 |
| ***otites*** | | 0,00 | 0,00 | 1,00E+00 | 0,00 | 0,00 | 7,41E-01 | **0,44** | **0,13** | **2,17E-146** | 0,09 | 0,13 | 7,14E-01 |
| ***pseudotites*** | | 0,00 | 0,00 | 9,99E-01 | 0,00 | 0,00 | 6,80E-01 | **0,46** | **0,16** | **3,81E-130** | 0,07 | 0,13 | 1,21E-01 |
| ***vulgaris*** | | 0,00 | 0,01 | 1,00E+00 | 0,00 | 0,00 | 1,00E+00 | **0,50** | **0,10** | **1,37E-117** | 0,03 | 0,07 | 9,99E-01 |

**<u>Supplementary Table S10:</u>** same as Supplementary Table S8, but only on autosomal genes.

| | | Expression increased compared to outgroup | | | | | | Expression decreased compared to outgroup | | | | | |
|---|---|---|---|---|---|---|---|---|---|---|---|---|---|
| | | Female ΔX | | | Male ΔX | | | Female ΔX | | | Male ΔX | | |
| Silene species | Sex-bias type | Proportion of sex-biased genes with high ΔX | Proportion of unbiased genes with high ΔX | corrected Chi2 p-value | Proportion of sex-biased genes with high ΔX | Proportion of unbiased genes with high ΔX | corrected Chi2 p-value | Proportion of sex-biased genes with high ΔX | Proportion of unbiased genes with high ΔX | corrected Chi2 p-value | Proportion of sex-biased genes with high ΔX | Proportion of unbiased genes with high ΔX | corrected Chi2 p-value |
| *acaulis* | Female-biased genes | 0,00 | 0,04 | 1,00E+00 | 0,00 | 0,03 | 1,00E+00 | NA | 0,41 | NA | 0,33 | 0,42 | 1,00E+00 |
| *colpophylla* | | 0,40 | 0,31 | 3,45E-01 | 0,09 | 0,14 | 6,54E-01 | 0,67 | 0,49 | 8,20E-01 | 0,62 | 0,35 | 1,53E-01 |
| *diclinis* | | 0,11 | 0,16 | 1,73E-01 | 0,10 | 0,14 | 8,49E-01 | 0,20 | 0,35 | 8,93E-02 | **0,54** | **0,35** | **1,92E-05** |
| *dioica* | | **0,28** | **0,18** | **2,45E-06** | 0,05 | 0,09 | 4,90E-01 | 0,21 | 0,32 | 1,68E-01 | **0,47** | **0,27** | **2,87E-13** |
| *heuffelli* | | 0,18 | 0,19 | 8,60E-01 | 0,10 | 0,10 | 1,00E+00 | **0,20** | **0,35** | **3,96E-02** | **0,43** | **0,31** | **5,07E-06** |
| *latifolia* | | **0,20** | **0,09** | **7,41E-09** | 0,14 | 0,21 | 3,91E-01 | 0,18 | 0,25 | 6,09E-01 | **0,58** | **0,39** | **9,35E-07** |
| *marizii* | | 0,17 | 0,15 | 7,68E-01 | 0,12 | 0,12 | 1,00E+00 | 0,23 | 0,35 | 6,18E-02 | **0,55** | **0,36** | **4,72E-09** |
| *nutans* | | 0,10 | 0,09 | 1,00E+00 | 0,00 | 0,03 | 7,68E-01 | 0,00 | 0,51 | 1,00E+00 | 0,35 | 0,29 | 8,93E-01 |
| *otites* | | 0,16 | 0,15 | 1,00E+00 | 0,06 | 0,12 | 8,60E-01 | 0,33 | 0,35 | 1,00E+00 | 0,56 | 0,37 | 6,54E-02 |
| *pseudotites* | | 0,19 | 0,15 | 6,09E-01 | 0,07 | 0,09 | 1,00E+00 | 0,42 | 0,41 | 1,00E+00 | **0,57** | **0,34** | **3,22E-03** |
| *vulgaris* | | 0,16 | 0,18 | 8,60E-01 | **0,02** | **0,11** | **1,34E-02** | 0,00 | 0,42 | 8,12E-01 | 0,35 | 0,42 | 5,28E-01 |
| *acaulis* | Male-biased genes | 0,04 | 0,04 | 1,00E+00 | 0,02 | 0,03 | 1,93E-01 | **0,65** | **0,41** | **1,54E-35** | **0,21** | **0,42** | **8,91E-05** |
| *colpophylla* | | **0,14** | **0,31** | **2,07E-03** | **0,04** | **0,14** | **1,27E-04** | **0,66** | **0,49** | **1,64E-04** | 0,28 | 0,35 | 6,09E-01 |
| *diclinis* | | **0,05** | **0,16** | **3,61E-02** | **0,03** | **0,14** | **2,63E-06** | **0,64** | **0,35** | **6,24E-21** | **0,03** | **0,35** | **2,59E-06** |
| *dioica* | | **0,06** | **0,18** | **4,88E-03** | **0,03** | **0,09** | **8,48E-05** | **0,68** | **0,32** | **1,04E-52** | **0,14** | **0,27** | **3,11E-02** |
| *heuffelli* | | **0,06** | **0,19** | **1,02E-03** | **0,02** | **0,10** | **2,58E-07** | **0,61** | **0,35** | **1,93E-21** | **0,17** | **0,31** | **3,85E-02** |
| *latifolia* | | 0,04 | 0,09 | 6,09E-01 | 0,17 | 0,21 | 1,48E-01 | **0,74** | **0,25** | **5,07E-98** | **0,21** | **0,39** | **1,95E-03** |
| *marizii* | | **0,07** | **0,15** | **4,07E-02** | **0,04** | **0,12** | **8,29E-07** | **0,66** | **0,35** | **1,44E-34** | **0,13** | **0,36** | **1,03E-05** |
| *nutans* | | 0,00 | 0,09 | 1,00E+00 | **0,01** | **0,03** | **9,95E-03** | **0,96** | **0,51** | **4,78E-80** | **0,14** | **0,29** | **1,24E-02** |
| *otites* | | 0,06 | 0,15 | 2,08E-01 | **0,03** | **0,12** | **2,07E-03** | **0,55** | **0,35** | **7,27E-06** | 0,27 | 0,37 | 8,62E-01 |
| *pseudotites* | | 0,06 | 0,15 | 1,02E-01 | **0,03** | **0,09** | **2,69E-03** | **0,62** | **0,41** | **1,58E-08** | 0,27 | 0,34 | 6,99E-01 |
| *vulgaris* | | **0,04** | **0,18** | **4,74E-03** | 0,08 | 0,11 | 1,73E-01 | **0,80** | **0,42** | **1,11E-43** | **0,13** | **0,42** | **7,68E-03** |

**Supplementary Table S11:** same as Supplementary Table S8 on **leaf** RNA-seq data.

| | | **Expression increased compared to outgroup** | | | | | | **Expression decreased compared to outgroup** | | | | | |
|---|---|---|---|---|---|---|---|---|---|---|---|---|---|
| | | **Female ΔX** | | | **Male ΔX** | | | **Female ΔX** | | | **Male ΔX** | | |
| **Silene species** | **Sex-bias type** | **Proportion of sex-biased genes with high ΔX** | **Proportion of unbiased genes with high ΔX** | **corrected Chi2 p-value** | **Proportion of sex-biased genes with high ΔX** | **Proportion of unbiased genes with high ΔX** | **corrected Chi2 p-value** | **Proportion of sex-biased genes with high ΔX** | **Proportion of unbiased genes with high ΔX** | **corrected Chi2 p-value** | **Proportion of sex-biased genes with high ΔX** | **Proportion of unbiased genes with high ΔX** | **corrected Chi2 p-value** |
| ***latifolia*** | **Female-biased genes** | 0,00 | 0,04 | 1,00 | 0,00 | 0,09 | 1,0000 | 0,00 | 0,36 | 1,0000 | 0,75 | 0,44 | 1,00 |
| ***latifolia*** | **Male-biased genes** | 0,00 | 0,04 | 1,00 | **0,33** | **0,09** | **0,0043** | **0,67** | **0,36** | **0,0256** | 0,50 | 0,44 | 1,00 |

**Supplementary Table S12:** Numbers of sex-linked genes that are female-biased or male-biased and for which the changes of expression are under selection or not in both sexes.

| **Species** | **# sex-linked female biased genes under selection** | **# sex-linked female biased genes not under selection** | **# sex-linked male biased genes under selection** | **# sex-linked male biased genes not under selection** | **P-value** |
|---|---|---|---|---|---|
| ***S. diclinis*** | 72 | 32 | 46 | 27 | $4.83e^{-01}$ |
| ***S. dioica*** | 107 | 22 | 39 | 35 | $\mathbf{8.48e^{-06}}$ |
| ***S. heuffelii*** | 118 | 7 | 22 | 22 | $\mathbf{8.84e^{-11}}$ |
| ***S. latifolia*** | 96 | 2 | 32 | 10 | $\mathbf{1.02e^{-04}}$ |
| ***S. marizii*** | 61 | 14 | 49 | 35 | $\mathbf{3.04e^{-03}}$ |
| ***S. colpophylla*** | 0 | 3 | 15 | 25 | $4.92e^{-01}$ |
| ***S. otites*** | 6 | 1 | 6 | 7 | $2.13e^{-01}$ |
| ***S. pseudotites*** | 2 | 0 | 9 | 7 | $6.69e^{-01}$ |

**Supplementary Table S13:** Numbers of sex-linked genes or autosomal genes that are female-biased and for which the changes of expression are under selection or not in both sexes.

| Species | # sex-linked female biased-genes under selection | # sex-linked female biased-genes | # autosomal female biased-genes under selection | # autosomal female biased-genes | P-value |
|---|---|---|---|---|---|
| ***S. diclinis*** | 72 | 32 | 107 | 83 | **$4.09e^{-02}$** |
| ***S. dioica*** | 107 | 22 | 265 | 143 | **$1.75e^{-04}$** |
| ***S. heuffelii*** | 118 | 7 | 248 | 210 | **$3.73e^{-16}$** |
| ***S. latifolia*** | 96 | 2 | 182 | 87 | **$4.79e^{-09}$** |
| ***S. marizii*** | 61 | 14 | 203 | 104 | **$1.57e^{-02}$** |
| ***S. colpophylla*** | 0 | 3 | 36 | 26 | $1.67e^{-01}$ |
| ***S. otites*** | 6 | 1 | 31 | 22 | $3.28e^{-01}$ |
| ***S. pseudotites*** | 2 | 0 | 51 | 30 | $7.40e^{-01}$ |

**Supplementary Table S14:** Generalized linear models (GLMs) between the number of sex-biased genes and the effective population size $N_e$ (the synonymous nucleotide diversity $\pi_S$ was used as a proxy). Please refer to Materials and Methods (M&M) equations 3 and 5 for details. See Supplementary Figure S2 for a graphical representation. None of the models were significant, showing there is no correlation between the number of sex-biased genes and the effective population size $N_e$.

| number of sex-biased genes | Model type | Estimate | Standard Error | Z-value | p-value |
|---|---|---|---|---|---|
| **Total** | M&M equation 3 (GLM, negative binomial family, offset) | -0.01483 | 0.15583 | -0.095 | 0.924 |
| **female-biased** | | 0.1842 | 0.2038 | 0.904 | 0.366 |
| **male-biased** | | -0.0781 | 0.14255 | -0.548 | 0.584 |
| **Total** | M&M Equation 5 (generalized least squares model correcting for phylogeny) | -0.00089 | 0.1911039 | -0.00467 | 0.9964 |
| **female-biased** | | -0.17501 | 0.2683476 | 0.652193 | 0.5351 |
| **male-biased** | | -0.05659 | 0.1775280 | -0.31878 | 0.7592 |

**Supplementary Table S15:** List of the four subsets of genes for which we tested an enrichment in functions or pathways, and the set of genes used as the background.

| | **Subset tested for an enrichment** | **"background" set of genes** |
|---|---|---|
| **(1)** | Whole male-biased or female-biased genes | Whole annotation |
| **(2)** | Male-biased or female-biased genes in gynodioecious species | Whole male-biased or female-biased genes |
| **(3)** | Male-biased or female-biased genes in each section of dioecious species | Male-biased or female-biased genes in the three sections of dioecious species |
| **(4)** | Male-biased or female-biased genes with a signature of positive selection | Whole male-biased or female-biased genes |

**Supplementary Table S16:** Number of individuals and tissues sampled for each species.

| Species name | Sexual system | Tissue | Number of individuals |
|---|---|---|---|
| ***Silene acaulis*** | Dioecious | Flower buds | 7 males and 8 females |
| ***Silene colpophylla*** | Dioecious | Flower buds | 7 males and 7 females |
| ***Silene diclinis*** | Dioecious | Flower buds | 6 males and 6 females |
| ***Silene dioica*** | Dioecious | Flower buds | 6 males and 6 females |
| ***Silene heuffelii*** | Dioecious | Flower buds | 6 males and 6 females |
| ***Silene latifolia*** | Dioecious | Flower buds | 5 males and 5 females |
| ***Silene latifolia*** | Dioecious | Leaves | 4 males and 4 females |
| ***Silene marizii*** | Dioecious | Flower buds | 6 males and 6 females |
| ***Silene nutans*** | Gynodioecious | Flower buds | 7 hermaphrodites and 4 females |
| ***Silene otites*** | Dioecious | Flower buds | 6 males and 6 females |
| ***Silene paradoxa*** | Hermaphrodite | Flower buds | 12 hermaphrodites |
| ***Silene pseudotites*** | Dioecious | Flower buds | 6 males and 6 females |
| ***Silene viscosa*** | Hermaphrodite | Flower buds | 3 hermaphrodites |
| ***Silene viscosa*** | Hermaphrodite | Leaves | 2 hermaphrodites |

| | | | |
|---|---|---|---|
| ***Silene vulgaris*** | Gynodioecious | Flower buds | 5 hermaphrodites and 4 females |
| ***Dianthus chinensis*** | Hermaphrodite | Seedlings | 2 hermaphrodites |
